# The USP28-ΔNp63 axis is a vulnerability of squamous tumours

**DOI:** 10.1101/683508

**Authors:** Cristian Prieto-Garcia, Oliver Hartmann, Michaela Reissland, Fabian Braun, Thomas Fischer, Susanne Walz, Annalena Fischer, Marco A. Calzado, Amir Orian, Mathias Rosenfeldt, Martin Eilers, Markus E. Diefenbacher

**Author notes:** Corresponding Author: Dr. Markus E. Diefenbacher. Lehrstuhl für Biochemie und Molekularbiologie, Biozentrum, Am Hubland, Würzburg, 97074, Germany.

## Abstract

The transcription factor ΔNp63 is a master regulator that establishes epithelial cell identity and is essential for the survival of SCC of lung, head and neck, oesophagus, cervix and skin. Here, we report that the deubiquitylase USP28 stabilizes ΔNp63 protein and maintains elevated ΔNP63 levels in SCC by counteracting its proteasome-mediated degradation. Interference with USP28 activity by genetic means abolishes the transcriptional identity of SCC cells and suppresses growth and survival of human SCC cells. CRISPR/Cas9-engineered mouse models establish that both induction and maintenance of lung SCC strictly depend on endogenous USP28. Targeting ΔNp63 protein abundance in SCC via inhibition of USP28 therefore is a feasible strategy for the treatment of SCC tumours.

**Significance:** SCC depend on ΔNp63, and its protein abundance is tightly controlled by the ubiquitin proteasome system. Here, we demonstrate the dependence of SCC on USP28 for various human SCC *in vitro* and *in vivo* using murine lung tumour models. As inhibitors for deubiquitylases become available, targeting USP28 is a promising therapeutic strategy.

## Introduction

Each year, around two million patients are diagnosed and approximately 1.76 million succumb to lung cancer, making this tumour entity by far the most leading cause of cancer related death, for men and women, worldwide (WHO cancer statistics, 2018). According to the current WHO classification lung cancer is classified into two major subtypes depending on marker expression: Non-Small Cell Lung Cancer (NSCLC) and Small Cell Lung Cancer (SCLC), causing around 85% or 15% of disease incidences, respectively (Inamura, 2017). NSCLC can be further subdivided according to marker expression and prevalent mutational aberrations, into adenocarcinomas (ADC) and squamous carcinomas (SCC). Comprehensive analyses of the mutational landscape show that lung SCC is one of the genetically most complex tumours (Cancer Genome Atlas Research, 2012). As consequence, little is known about therapeutic targetable vulnerabilities of this disease.

A key regulatory protein in SCC is the p53-related transcription factor ΔNp63, encoded by the *TP63* gene (Su et al., 2013). ΔNp63 is highly expressed in lung SCC as well as in SCCs of the skin, head and neck, and oesophagus, in part due to gene amplification (Cancer Genome Atlas Research, 2012; Hibi et al., 2000; Tonon et al., 2005). The *TP63* locus encodes multiple mRNAs that give rise to functionally distinct proteins. Notably, transcription from two different promoters produces N-terminal variants either containing or lacking the transactivation domain: TAp63 or ΔNp63 (Deyoung and Ellisen, 2007). The major p63 isoform expressed in squamous epithelium and SCC is ΔNp63α(Koster et al., 2007; Rocco et al., 2006), which is a master transcription factor that establishes epithelial cell identity, including cytokeratin 5/6 and 14 (Deyoung and Ellisen, 2007; Hamdan and Johnsen, 2018; Rocco et al., 2006; Somerville et al., 2018; Su et al., 2013). In addition, ΔNp63 binds to and thereby inactivates TP53 at promoters of pro-apoptotic genes, suppressing their expression (Craig et al., 2010; Westfall et al., 2003). ΔNp63 is essential for the survival of skin and pancreatic SCC cells, since established murine skin SCCs are exquisitely dependent on ΔNp63; acute deletion of *TP63* in advanced, invasive SCC induced rapid and dramatic apoptosis and tumour regression (Galli et al., 2010; Ramsey et al., 2013; Rocco et al., 2006; Somerville et al., 2018; Su et al., 2013). Collectively, these findings raise the possibility that ΔNp63 is a therapeutic target in SCC tumours.

ΔNp63 is an unstable protein that is continuously turned over by the proteasome upon ubiquitination by E3-ligases, such as the FBXW7 ubiquitin ligase (Galli et al., 2010). *FBXW7* is frequently mutated in SCC tumours (Galli et al., 2010; Ruiz et al., 2019). Intriguingly, it has been shown previously that the degradation of many targets of FBXW7 is counteracted by the deubiquitylase (DUB) USP28 (Popov et al., 2007b). This is in part due to the fact that USP28 exploits binding to FBXW7 to interact with its substrates (Schulein-Volk et al., 2014). However, USP28 can also recognize the phosphodegron, required for the binding of FBXW7 to its substrates in an FBXW7-independent manner (Diefenbacher et al., 2015). Loss of USP28 counteracts the loss of Fbxw7 in a murine colon tumour model (Cremona et al., 2016; Diefenbacher et al., 2015), and acute deletion of USP28 in established tumours increases survival in the APC^minΔ/+^ colorectal tumour model (Diefenbacher et al., 2014), while not affecting tissue homeostasis in non-transformed cells (Schulein-Volk et al., 2014). Together, these data argue that targeting USP28 may destabilize ΔNp63 and suggest that this strategy may have therapeutic efficacy in SCC.

## Results

### USP28 is highly abundant in human squamous tumours and correlates with poor prognosis

To investigate the mutational as well as the expression status of USP28 in lung cancer, we analysed publicly available datasets of human tumours (Figure 1A, B and Figure S1A, B, D). USP28 is rarely lost nor mutated, but frequently transcriptionally upregulated in human SCC compared to normal tissue or ADC (adenocarcinoma) patient samples (Figure 1A, B and S1A, B). Similarly, expression of *TP63* was significantly upregulated in SCC samples compared to non-transformed tissue or to ADC samples (Figure 1A and Figure S1A, B).

**Figure 1:**
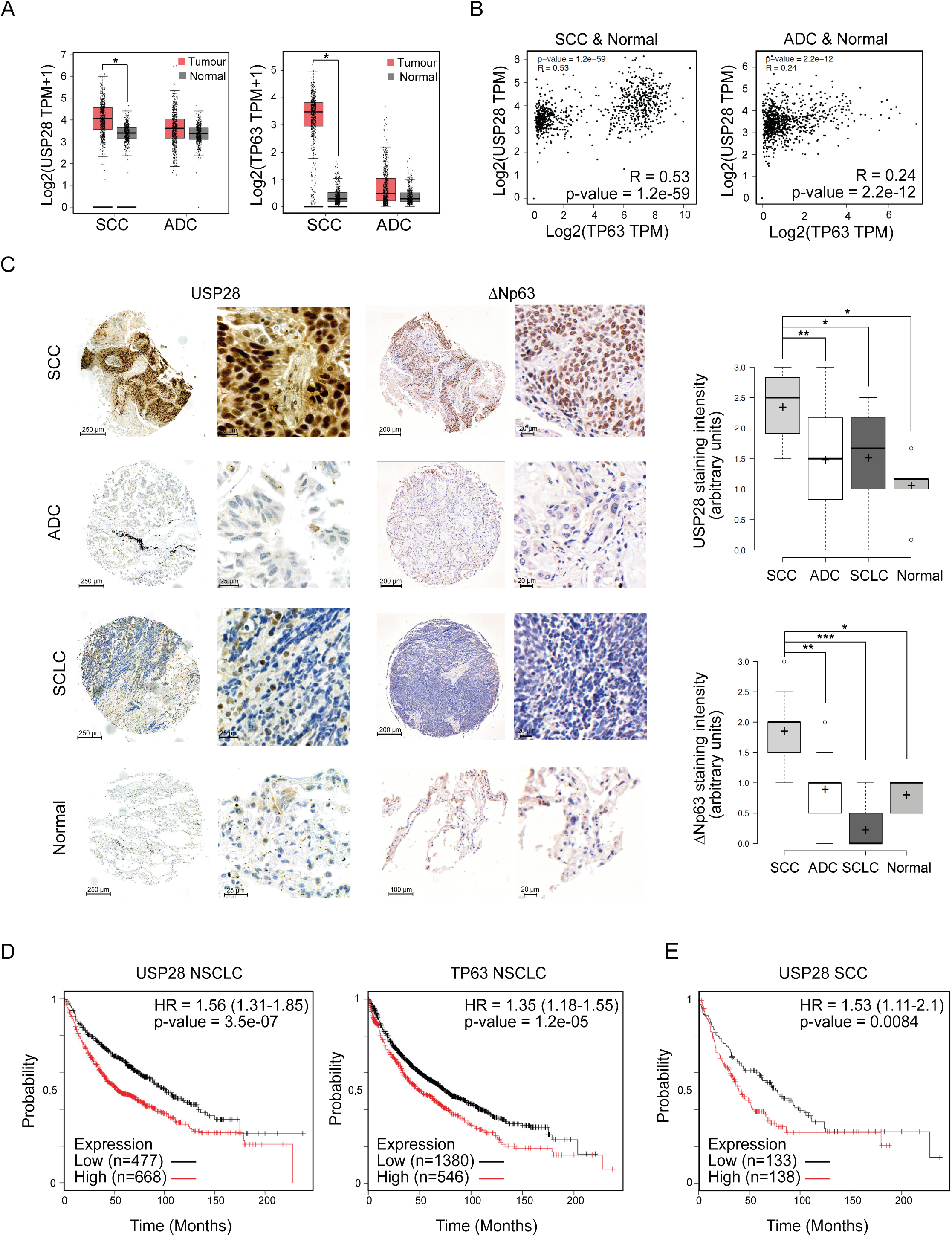
USP28 is highly abundant in human squamous tumours and correlates with poor prognosis. A. Expression of USP28 (left) and TP63 (righ) in human lung squamous cell carcinomas (SCC, n=486), adenocarcinomas (ADC, n=483) and normal non-transformed tissue (normal SCC=50, normal ADC=59). p-values were calculated with a one-way ANOVA comparing normal tissue tumours. TPM: transcript count per million mapped reads B. Correlation of mRNA expression of USP28 and TP63 in lung SCC (left, n=824), ADC (right, n=830) and normal, non-transformed tissue. R: Spearmans correlation coefficient C. IHC analysis of USP28 and ΔNp63 protein abundance in lung cancer and non-transformed human samples (n=300). The staining intensity was quantified in arbitrary units from 0 up to 3 (Figure S1B) by three independent pathologists. In box plots, the centre line reflects the median, the cross represents the mean and the upper and lower box limits indicates the first and third quartile. Whiskers extend 1.5x the IQR and outliers are marked as dots. p-values were calculated using two-tailed t-test. D. Kaplan-Meier estimator of NSCLC patients stratified by USP28 (left, n=1145) and TP63 (right, n=1926) expression. p-values were calculated using log-rank test. HR: hazard ratio E. Kaplan-Meier estimator of lung SCC patients stratified by USP28 expression (n=271). The p-value was calculated using a logrank test. HR: hazard ratio *p-value < 0.05; **p-value < 0.01; *** p-value < 0.001; see also Figure S1.

Next, we determined the abundance of USP28 protein via immuno-histochemistry (IHC) on tissue microarrays and tumour sections of a total of 300 human lung tumour samples. Relative to non-transformed tissue, all samples from different human lung tumour subtypes expressed elevated levels of USP28, with SCC presenting the highest levels (Figure 1C), confirming the USP28 mRNA expression data (Figure 1A). TP63 protein abundance was evaluated within the same cohort and, like USP28, exhibited the highest protein abundance in SCC tumours, when compared to ADC and SCLC samples and normal tissue (Figure 1C). To evaluate the relevance of both proteins for tumour development, we used publicly available datasets to correlate mRNA expression data with patient survival. Patients with an increased expression of either Δ*Np63* or *USP28* showed a significantly shortened overall survival (Figure 1D). Importantly, this correlation was not a secondary consequence of a generally shorter survival of SCC patients, since USP28 expression correlated with worse prognosis even when only SCC patients were analysed (Figure 1E). Finally, we noted that 3% of lung SCC patients display mutations in *USP28* or a deletion of *USP28,* and those showed a much better disease-free survival compared to USP28 wildtype patients (Figure S1D). These data indicate that USP28 is upregulated in NSCLC, and high expression of USP28 negatively correlates with overall patient survival in SCC tumours. Additionally, we were able to detect a strong correlation between USP28 and ΔNp63 abundance in lung SCC, indicating a potential crosstalk between both proteins.

### ΔNp63 stability is regulated by USP28 via its catalytic activity

To test whether USP28 controls ΔNp63 protein abundance, we initially expressed HA-tagged UPS28 and FLAG-tagged ΔNp63 in HEK293 cells by transient transfection. Immunofluorescence staining using antibodies against USP28 and ΔNp63 revealed that both proteins localise to the nucleus of transfected cells (Figure S2A). Co-immunoprecipitation experiments showed that ΔNp63 binds to USP28 and *vice versa*, indicating an interaction of both molecules in cells (Figure 2A). Upon co-expression of His-tagged ubiquitin, ΔNp63 was ubiquitylated, as demonstrated by pulldown of His-tagged ubiquitin, followed by immunoblot using a ΔNp63 specific antibody (Figure 2B) and co-expression of USP28 resulted in the de-ubiquitylation of ΔNp63 (Figure 2B). To test the chain specificity of substrate de-ubiquitylation by USP28 on ΔNp63, we ectopically co-expressed a His-tagged ubiquitin that carries a single lysine residue either at position K48 or K63. Upon His-ubiquitin pulldown, K48-as well as K63-linked poly-ubiquitin chains could be detected on ΔNp63, as previously reported (Galli et al., 2010; Peschiaroli et al., 2010) (Figure 2C). Upon overexpression of USP28, only K48-linked ubiquitin chains were removed from ΔNp63, whereas K63-linked chains were resistant to USP28 (Figure 2C). To test whether USP28 catalytic activity is required for de-ubiquitination of ΔNp63, we used a catalytic inactive mutant of USP28, USP28^C171A^ (Figure 2D-F) (Diefenbacher et al., 2015; Diefenbacher et al., 2014; Popov et al., 2007b; Schulein-Volk et al., 2014). Immunoprecipitation of transfected cells using an ΔNp63-specific antibody revealed that USP28^C171A^ was able to bind to ΔNp63 (Figure 2D). While overexpression of the wild type form of USP28 deubiquitylated ΔNp63 (Figure 2E), USP28^C171A^ failed to do so, demonstrating that the catalytically active cysteine of USP28 is required for deubiquitylation of ΔNp63 (Figure 2E).

**Figure 2:**
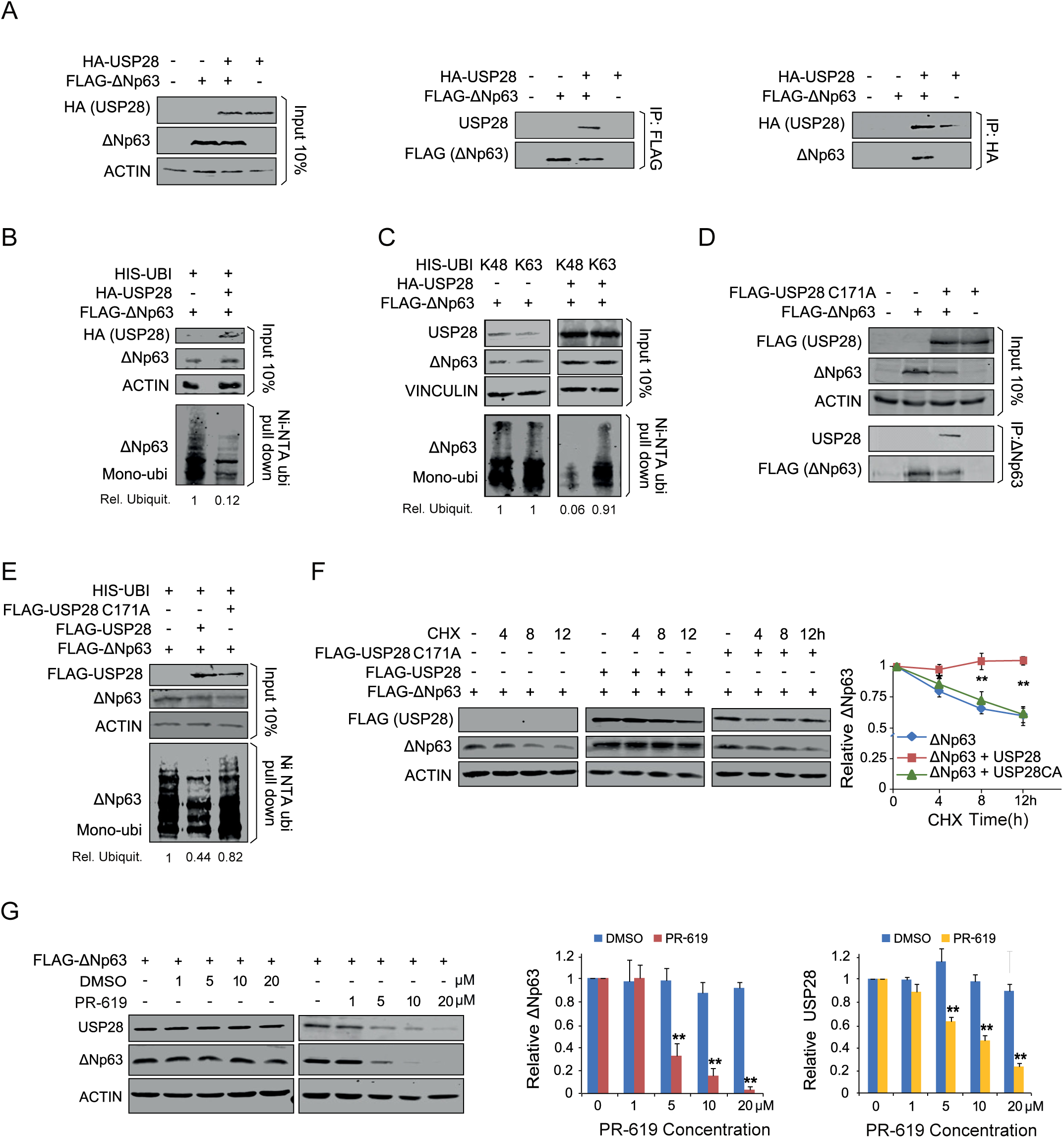
ΔNp63 stability is regulated by USP28 via its catalytic activity. A. Co-immunoprecipitation of exogenous HA-USP28 and FLAG-ΔNp63 in HEK293 cells. Either HA-USP28 or FLAG-ΔNp63 were precipitated and blotted against FLAG-ΔNp63 or HA-USP28, The input corresponds to 10% of the total protein amount used for the IP (ACTIN as loading control). B. Ni-NTA His-ubiquitin pulldown in control transfected or HA-USP28 overexpressing HEK293 cells, followed by immunoblot against exogenous ΔNp63. The input corresponds to 10% of the total protein amount used for the pull down. Relative ubiquitination was calculated using ACTIN for normalization. C. Ni-NTA His-ubiquitin pulldown K48 or K63 in control and HA-USP28 overexpressing HEK293 cells, followed by immunoblot against exogenous ΔNp63. The input corresponds to 10% of the total protein amount used for the pulldown. Relative ubiquitination was calculated using ACTIN for normalization. D. Co-immunoprecipitation of exogenous FLAG-USP28 C171A and FLAG-ΔNp63 in HEK293 cells. ΔNp63 was precipitated and blotted against FLAG-USP28 or ΔNp63. The input corresponds to 10% of the total protein amount used for the IP (ACTIN as loading control). E. Ni-NTA His-ubiquitin pulldown in control, FLAG-USP28 or FLAG-USP28 C171A transfected HEK293 cells, followed by immunoblot against exogenous ΔNp63. The input corresponds to 10% of the total protein amount used for the pulldown. Relative ubiquitination was calculated using ACTIN for normalization. F. CHX chase assay (100ug/ml) of control, FLAG-USP28 or FLAG-USP28 C171A transfected HEK293 cells for indicated time points. Representative immunoblot analysis of FLAG (USP28) and ΔNp63 as well quantification of relative protein abundance (ACTIN as loading control). G. Immunoblot of USP28 and ΔNp63 in transfected HEK293 cells upon treatment with either DMSO or indicated concentrations of PR-619 for 24 hours. Relative protein abundance was calculated ACTIN as loading control. Quantification of n=3 experiments, Significance was calculated using two-tailed t-test. All quantitative data are represented as mean ± SD (n=3). p-values were calculated using two-tailed t-test statistical analysis; *p-value < 0.05, **p-value < 0.01, *** p-value < 0.001; see also Figure S2.

K48-linked ubiquitin chains target proteins to the proteasome for degradation (Grice and Nathan, 2016). Since USP28 is able to counteract K48 linked ubiquitylation of ΔNp63, we investigated the ability of USP28 to modulate ΔNp63 protein turnover. To do so, we co-expressed ΔNp63 with either wildtype USP28 or USP28^C171A^ in HEK293 cells. 24 hours post transfection, cells were treated with 100 μg/ml cycloheximide (CHX) to block protein synthesis. ΔNp63 was degraded in control cells with a half-life of 8 hrs. Co-expression of wildtype USP28, but not of the catalytically inactive mutant, strongly stabilised ΔNp63 protein (Figure 2F).

As ΔNp63 protein stability was enhanced by USP28, but not the catalytic inactive C171A mutant, we tested whether a pharmacologic inhibitor of DUBs, PR-619, would also affect overall protein abundance of ΔNp63. Therefore, we expressed ΔNp63 in HEK293 cells and 24 hours post transfection treated cells with either DMSO or increasing amounts of PR-619 for additional 24 hours (Figure 2G). While ΔNp63 was not degraded in control cells treated with DMSO, cells exposed to PR-619 showed a shortened half-life of ΔNp63 and an IC_50_ of <5μM. with a half-life of 8 hrs. In control-treated cells, the protein abundance of USP28 was not affected, however, upon addition of the pan-DUB inhibitor PR-619, USP28 protein was reduced in a dose-dependent fashion (Figure 2G). This is in line with previous observations that the enzymatic activity of DUBs is required to enhance their own stability (de Bie and Ciechanover, 2011; Wang et al., 2017).

Collectively, these data demonstrate that USP28 can interact with and stabilize the ΔNp63 protein by removing K48-linked ubiquitin chains, and that the catalytic domain of USP28 is required for this activity.

### USP28 stabilizes ***Δ***Np63 independently of FBXW7

Previous reports highlighted the regulation of ΔNp63 protein stability by the E3-ligase FBXW7 (Galli et al., 2010). To investigate if USP28 interacts with ΔNp63 in a FBXW7-dependent fashion, we made use of a ΔNp63 point mutant, ΔNp63^S383A^, which is not phosphorylated by GSK3β and abolishes binding to FBXW7 (Galli et al., 2010). In transient expression assays using HEK293 cells we found that USP28 was able to bind to ΔNp63^S383A^ (Figure S2B and C). This interaction resulted in a decreased ubiquitylation of ΔNp63^S383A^ (Figure S2D). Furthermore, overexpression of USP28 was able to increase protein half-life (Figure S2E) and treatment of cells with PR-619 affected ΔNp63^S383A^ protein stability, albeit to a somewhat lesser extent compared to wild type ΔNp63 (Figure S2F).

To determine whether endogenous USP28 regulates the abundance and stability of ΔNp63, we used a human SCC cell line (A431). Both proteins were readily detectable in the nucleus of these cells by immunofluorescence (Figure 3A). Furthermore, immunoprecipitation of endogenous USP28 co-immunoprecipitated endogenous ΔNp63, and vice versa (Figure 3B). In contrast, antibodies against USP25, a ubiquitin-specific protease that is structurally very similar to USP28 (Gersch et al., 2019; Sauer et al., 2019), did not co-immunoprecipitate ΔNp63 although USP25 is readily detectable in A431 cells. Conversely, antibodies against ΔNp63 did not co-immunoprecipitate endogenous USP25 (Figure 3B). To investigate whether USP28 regulates ΔNp63 protein stability, we generated cell lines expressing either a doxycycline-inducible or constitutive shRNA targeting *USP28* (Figure 3C and S4C).

**Figure 3:**
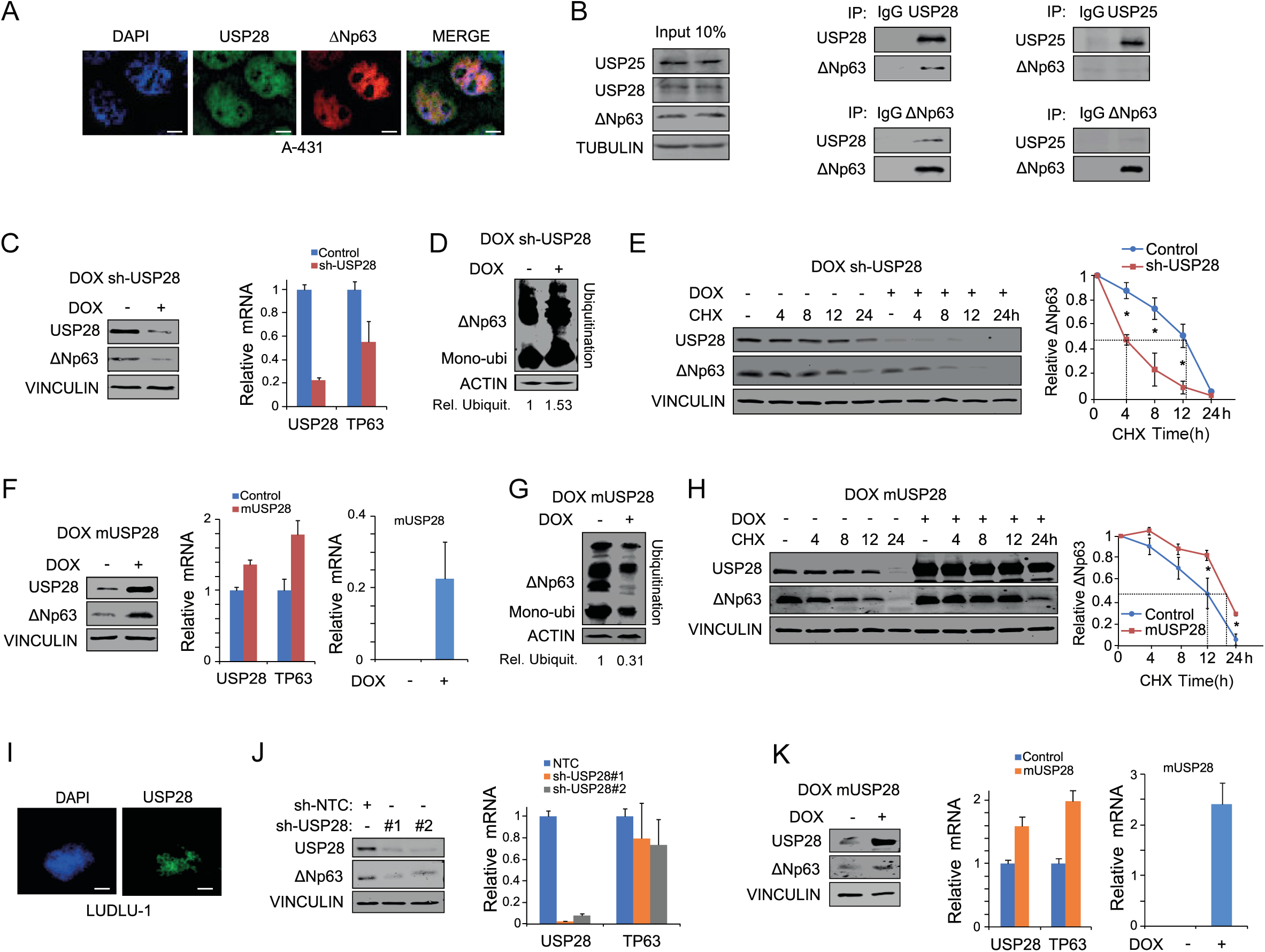
USP28 regulates ΔNp63 protein stability in SCC tumour cell lines. A. Immunofluorescence (IF) staining against endogenous USP28 and ΔNp63 in the SCC cell line A-431 (blue: DAPI [staining of nuclei], green/cyan: USP28 and red/magenta: ΔNp63), Scale bar = 5µm B. Immunoblot of endogenous USP25, USP28 and ΔNp63 immunoprecipitated (IP) from A-431 cells and co-precipitated ΔNp63 or USP28. Beads were coupled to specific USP25, USP28 and ΔNp63 antibodies or non-specific rabbit IgG as control. The input corresponds to 10% of the total protein amount used for the IP (TUBULIN as loading control). C. Inducible depletion of USP28 in A-431 upon treatment with doxycycline (1ug/ml) for 96h, western blot (left, VINCULIN as loading control) and qPCR analysis of USP28 and ΔNp63 expression relative to ACTIN (right) was performed. D. Tandem ubiquitin binding entity (TUBE) pulldown of endogenous ubiquitylated ΔNp63 in A-431 cells upon DOX depletion of USP28. Relative ubiquitination was calculated using ACTIN for normalization. E. Cycloheximide (CHX) chase assay (100ug/ml) of control or inducible sh-USP28 A-431 cell line (EtOH or 1ug/ml dox) for indicated time points. Representative immunoblot (left, VINCULIN as loading control) of USP28 and ΔNp63 and quantification of relative protein abundance (right). F. Doxycycline induced murine USP28 overexpression (EtOH or 1ug/ml dox for 96 hours) in A-431 cells followed by immunoblot (VINCULIN as loading control) and qPCR analysis of USP28 and ΔNp63. For qPCR, human USP28 and murine USP28 (mUSP28) primers were used. Relative mRNA was calculated using ΔΔCt analysis for human USP28 and ΔCt for mUSP28 (ACTIN as housekeeping). G. TUBE pulldown of endogenous ubiquitylated ΔNp63 in A431 cells upon overexpression of mUSP28 for 96 hours (EtOH or 1ug/ml dox). Relative ubiquitination was calculated using ACTIN for normalization. H. CHX chase assay (100ug/ml) of control or inducible mUSP28 overexpressing A-431 cell line (EtOH or 1ug/ml dox) for indicated time points. Representative immunoblot analysis of USP28 and ΔNp63 as well quantification of relative protein abundance (VINCULIN as loading control). I. IF staining against endogenous USP28 in human lung SCC cell line LUDLU-1 (blue: DAPI, green/cyan: USP28). Scale bar in µm J. Immunoblot of control (sh-NTC) and two independent shRNA targeting USP28 (sh-USP28#1 and #2) for ΔNp63 and USP28 protein abundance in LUDLU-1 (VINCULIN as loading control), followed by qPCR analysis of USP28 and ΔNp63 expression relative to ACTIN. K. Doxycycline induced mUSP28 overexpression (EtOH or 1ug/ml dox for 96 hours) in LUDLU-1 cells followed by immunoblot (VINCULIN as loading control) and qPCR analysis of USP28, mUSP28 and ΔNp63. Relative mRNA was calculated using ΔΔCt analysis for human USP28 and ΔCt for mUSP28 (ACTIN as housekeeping for the analysis). All quantitative data are represented as mean ± SD (n=3). p-values were calculated using two-tailed t-test statistical analysis; *p-value < 0.05; **p-value < 0.01; *** p-value < 0.001; see also Figure S3.

Firstly, we investigated the effects of acute USP28 depletion on ΔNp63 protein abundance using doxycycline inducible shRNA cell lines. ΔNp63 protein abundance was reduced in USP28 acutely depleted cells (Figure 3C) as detected by western blot. Notably, depletion of USP28 also reduced ΔNp63 mRNA levels, consistent with previous observations that ΔNp63 activates expression of its own mRNA (Antonini et al., 2006)(Figure 3C). To assess the impact of acute USP28 depletion on endogenous ΔNp63 ubiquitylation, we performed tandem <u>u</u>biquitin <u>b</u>inding <u>e</u>ntity (TUBE) pull-down assays (Hjerpe et al., 2009). TUBEs are composed of four copies of the ubiquitin-associated domain of Ubiquilin fused in tandem to a glutathione-S-transferase (GST) tag and enable the detection of endogenous ubiquitin modifications on target proteins. TUBE ubiquitin pulldown experiments of cell lines expressing inducible shRNA targeting USP28 revealed increased ubiquitylation of ΔNp63 upon reduction of USP28 (Figure 3D). Next, we asked whether acute loss of USP28 affects ΔNp63 protein half-life. Inducible shRNA cell lines were cultured in the presence of EtOH (control) or doxycycline (1µg/ml) for 24hours prior to CHX treatment for the indicated time points, and USP28 and ΔNp63 protein abundance measured by western blot. Acute loss of USP28 reduced ΔNp63 protein abundance to 50% within 4 hours, compared to a half-life of 12 hours observed in control cells (Figure 3E).

To test whether increasing the levels of USP28 is able to increase ΔNp63 stability in a SCC cell line, we generated a doxycycline inducible system for the overexpression of murine USP28 in A431 cells (Figure 3F-H). Upon treatment with doxycycline, elevated amounts of USP28 resulted in an increase in ΔNp63 protein levels (Figure 3F). The effects on ΔNp63 were not only detectable on protein level, but were also reflected in increased levels of ΔNp63 mRNA, most likely due to the positive autoregulation of p63 transcription discussed before (Figure 2F). Exogenous USP28 also resulted in the decreased ubiquitylation of ΔNp63 as detected by TUBE assay (Figure 3G) and enhanced protein stability, extending its half-life from 12 hours to around 20 hours (Figure 3H). This demonstrates the ability of USP28 to deubiquitylate ΔNp63 and stabilise the protein.

Indeed, A431 cells are homozygous for the *S462Y* mutation in FBXW7, which is thought to inactivate substrate recognition (Figure S3C and D) (Yeh et al., 2016). The observed regulation of ΔNp63 in A431 cells argues that USP28 stabilizes ΔNp63 in an FBXW7-independent manner. To confirm that USP28 can also regulate ΔNp63 in FBXW7 wild-type SCC cells, we depleted USP28 using two independent shRNAs in LUDLU-1 cells (Figure 3J). Similar to the results obtained in A431 cells, depletion of USP28 resulted in the reduction of ΔNp63 protein abundance (Figure 3J). Furthermore, doxycycline-inducible overexpression of murine USP28 further increased endogenous ΔNp63 protein levels (Figure 3K) and enhanced ΔNp63 mRNA levels, highlighting a positive feedback loop for ΔNp63. These data demonstrate that USP28 de-ubiquitinates and stabilizes ΔNp63 and that both, substrate recognition and stabilisation occur in an FBXW7 independent manner.

### The ΔNp63-USP28 axis is required to maintain the identity of SCC cells

Both SCC cells and tumours depend on ΔNp63 for maintaining proliferation and cell identity (Lau et al., 2013; McDade et al., 2011; Ramsey et al., 2013). To explore whether USP28 controls these biological functions of ΔNp63, we first targeted ΔNp63 by two independent shRNA sequences, and analysed the knock down efficacy by immunoblotting (Figure S4A). Both shRNAs led to a significant decrease in ΔNp63 protein levels (Figure S4A), while they had no effect on USP28 abundance. Control or ΔNp63-depleted A431 cells were seeded at equal cell density and counted at indicated time points (Figure S4B). Depletion of ΔNp63 decreased SCC (Figure S4B) and cell cycle profiling indicated a mild G1-S-phase arrest (Figure S4C), consistent with previous reports(Wang et al., 2019). Depletion of USP28 using two independent shRNAs, which decreased ΔNp63 levels by at least 70%, caused a very similar decrease in cell proliferation and cell cycle progression (Figure S4D to F).

To investigate if USP28, like ΔNp63, is required to maintain the characteristic gene expression pattern of SCC cells, RNA expression profiles of A431 cells stably expressing shRNAs targeting either USP28 or ΔNp63 were compared to cells expressing a non-targeting control shRNA. Analysis of global gene expression revealed a strong correlation of target gene regulation in response to depletion of USP28 or ΔNp63 (R=0.48, m=0.72, p-value <2.2^308, Figure 4A). Specifically, there were 266 commonly downregulated genes and 66 commonly upregulated genes (Figure 4B and C). Gene set enrichment analysis (GSEA) using genes downregulated in response to depletion of either USP28 or ΔNp63, respectively, confirmed the strong similarity in expression changes caused by depletion of either factor (Figure. 4D and E).

**Figure 4:**
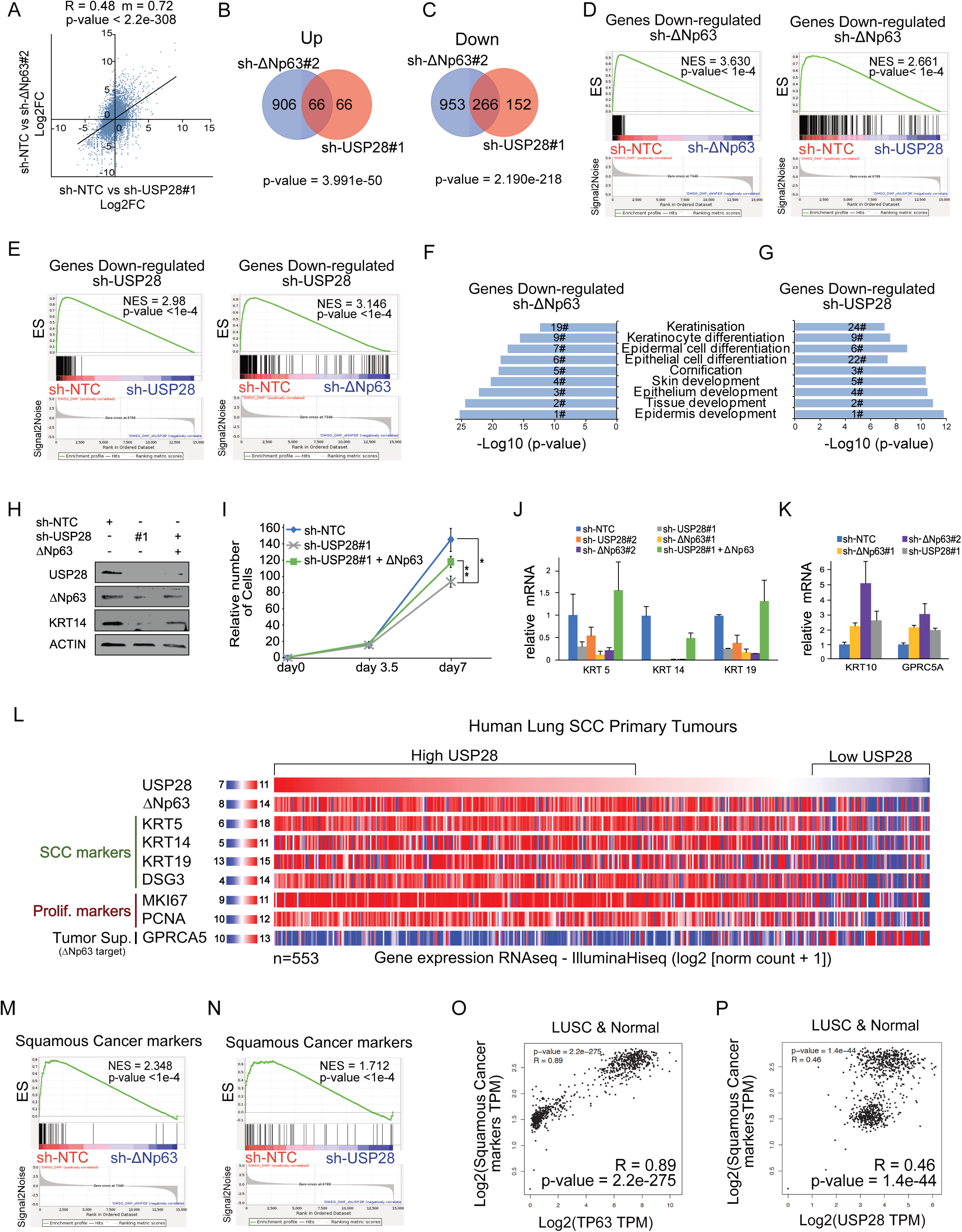
SCC tumour cells are dependent on USP28 and/or ΔNp63 to maintain a SCC identity. A. Correlation of gene expression changes upon constitutive transduction of A-431 cells with either shRNA targeting USP28 (sh-USP28#1) or ΔNp63 (sh-ΔNp63#2) relative to non-targeting control (sh-NTC). The diagonal line reflects a regression build on a linear model. R: Pearsons correlation coefficent, m: slope of the linear regression model B. Venn diagram of differentially up-regulated genes (log_2_FC>1.5 and q-value<0.05) between sh-ΔNp63#2 and sh-USP28#1 relative to non-targeting control (sh-NTC) in A-431 cells. p-values were calculated using a hypergeometric test. C. Venn diagram of differentially down-regulated genes (log_2_FC<-1.5 and q-value<0.05) between sh-ΔNp63#2 and sh-USP28#1 relative to sh-NTC in A-431 cells. p-values were calculated using a hypergeometric test. D. Gene set enrichment analysis (GSEA) of a gene set of significantly down-regulated genes in sh-ΔNp63#2 transfected A-431 cells (“Down-regulated sh-ΔNP63”, Supp. Table 1). The gene set was analysed in sh-ΔNp63#2-(left) and sh-USP28#1-depleted (right) A-431 cells. (N)ES: (normalised) enrichment score E. Gene set enrichment analysis (GSEA) of a gene set of significantly down-regulated genes in sh-USP28#1 transfected A-431 cells (“Down-regulated sh-USP28”, Supp. Table 1). The gene set was analysed in shUSP28#1-(left) and sh-ΔNp63#-depleted (right) A-431 cells. (N)ES: (normalised) enrichment score F. GO term analysis of biological processes enriched in sh-ΔNp63#2-depleted A-431 cells relative to sh-NTC. Numbers indicate the ranking position of all analysed GO terms based on the significance. G. GO term analysis of biological processes enriched in sh-USP28#1-depleted A-431 cells relative to sh-NTC. Numbers indicate the ranking position of all analysed GO terms based on the significance. H. Immunoblot of endogenous ΔNp63 and USP28 in A431 cells stably transduced with constitutive shRNA-non-targeting control (NTC) or against USP28 and transiently transfected with exogenous ΔNp63. Actin served as loading control. I. Cell growth of A431 cells stably transduced with constitutive shRNA-non-targeting control (NTC) or against USP28 and transiently transfected with exogenous ΔNp63. Total cell number was measured in triplicate and assessed at indicated time points. *p-value < 0.05, **p-value < 0.01 J. Relative expression of SCC marker genes KRT5, 14 and 19 in A431 cells stably transduced with constitutive shRNA-non-targeting control (NTC), two independent shRNA-ΔNp63, two independent shRNA-USP28 or ΔNp63 in shRNA-USP28#1, normalised to ACTIN. K. Relative expression of epithelial marker genes KRT10 and GPCR5A in A431 cells stably transduced with constitutive shRNA-non-targeting control (NTC), two independent shRNA-ΔNp63 and shRNA-USP28#1, normalised to ACTIN. L. Genomic signature of primary human lung SCC samples comprising USP28, ΔNp63, KRT5, KRT14, KRT19, DSG3, MKI67, PCNA and GPRC5A. Samples were sorted dependent on relative USP28 expression (high to low). n=553. M. GSEA of consensus squamous cancer marker genes (see Supp. Table 2) in A-431 cells stably transduced with sh-ΔNp63#2 or sh-NTC. (N)ES: (normalised) enrichment score N. GSEA of consensus squamous cancer marker genes in A-431 cells stably transduced with sh-USP28#1 or sh-NTC. (N)ES: (normalised) enrichment score O. Correlation of mRNA expression of consensus squamous cancer marker genes and TP63 in lung SCC and non-transformed lung tissue (Normal). R: Spearmans correlation coefficent. P. Correlation of mRNA expression of consensus squamous cancer marker genes and USP28 in lung SCC and non-transformed lung tissue (Normal). R: Spearmans correlation coefficent See also Figure S4.

To gain insight into the biological processes that underlie this similarity, we performed GO-term analysis and found that depletion of either ΔNp63 or USP28 strongly downregulates genes mapping to a set of overlapping GO-terms that describe epithelial cell identity and keratin expression (Figure 4F and G). As SCC depend on the expression of ΔNp63 to maintain proliferation (Figure S4A and B), we wondered if overexpression of ΔNp63 is able to rescue the observed proliferative defect in USP28 knock-down A431 cells. While depletion of USP28 reduced ΔNp63 and KRT14 protein abundance, exogenous ΔNp63 restored KRT14 expression in USP28 knock down cells, and partially restored proliferation (Figure 4H and I). This data indicates that depletion of USP28 affected cellular proliferation in SCC via reducing ΔNp63 levels.

To determine whether USP28 regulates genes involved in epithelial cell identity via ΔNp63, we tested whether ectopic expression of ΔNp63 restores the expression of these genes in USP28-depleted cells. RT-PCR analyses showed that depletion of ΔNp63 or USP28 decreases expression of keratins KRT5, KRT14 and KRT19 in A431 cells and expression of exogenous ΔNp63 partially restored expression of these genes in USP28-depleted cells (Figure 4J), demonstrating that the effect of USP28 is mediated in part by the downregulation of ΔNp63. In contrast to SCC associated cytokeratins, a differentiation associated marker, KRT10(Saladi et al., 2017), was upregulated in ΔNp63 and USP28 depleted A431. Similar responses were observed for the putative tumour suppressor GPRC5A(Saladi et al., 2017); loss of either protein resulted in an increased expression. In analysing publicly available RNA-Sequencing data of primary human lung SCC we could observe a similar correlation between USP28, ΔNp63 and SCC marker gene expression, such as KRT5, 14 and 19 and the tumour suppressor GPRC5A (Figure 4L).

Since the transcriptional effects observed in the knock-down cell lines by GO analysis hinted towards pathways frequently found deregulated in SCC tumours (Koster et al., 2007; Rocco et al., 2006), which was also observed by the strong effects on KRT5, KRT14 and KRT19 expression (Figure 4J, K and L), we next wanted to know if other SCC markers are also regulated by the USP28/ΔNp63 axis. Therefore, we analysed a panel of SCC relevant marker genes, which have previously been shown to be expressed in all SCC subclasses (Ferone et al., 2016; Mukhopadhyay et al., 2014; Wilkerson et al., 2010; Xu et al., 2014) (Figure 4L and Supplementary table 2). We analysed the expression of these genes in control, ΔNp63 and USP28 knock-down A431 cells. In cells depleted of ΔNp63, genes associated with SCC were downregulated, in line with the function of ΔNp63 as a master regulator of SCC (Figure 4L). Analysing the SCC marker gene panel in USP28 knock down cells also revealed a striking similarity to ΔNp63 knock down (Figure 4M). Furthermore, SCC markers were commonly downregulated in cells with reduced amounts of USP28 (Fig. 4M and Figure S4G to I). To assess if the observed genetic alterations are attributed to the USP28-ΔNp63 axis in SCC, we next analysed ‘Hallmark’ gene sets of reported USP28 substrates NOTCH1, MYC and AP1-cJUN, in USP28 and ΔNp63 knock down A431 cells (Figure SJ and K). In neither case a significant downregulation, as seen for ΔNp63 (Figure 4D), could be observed, indicating that the biological effects observed in SCC are specific to the regulation of ΔNp63 by USP28. Additionally, no compensatory gene expression mechanism was observed in ΔNp63 silenced cells (Figure S4K), demonstrating that ΔNp63 is a vulnerability of SCC.

Furthermore, publicly available expression datasets highlight the strong correlation of ΔNp63 and SCC marker co-expression in human tumour samples (Spearman R=0.89, p=2,2e-275; Figure 4N) and in line with the observed dependency of SCC cells for USP28, a correlation between SCC marker expression and USP28 abundance (Spearman R=0,46, p=1,4e-44; Fig. 4O).

In conclusion, targeting ΔNp63 or USP28 revealed that both factors are implicated in the regulation of a highly similar transcriptional network, in particular controlling the expression of SCC marker genes as well as genes controlling cell fate and SCC identity.

### Targeting USP28 during tumour initiation blocks SCC formation

Since available USP28 inhibitors (Wrigley et al., 2017) are not yet suited for *in vivo* application, we used genetic tools to interrogate the role of USP28 in induction and maintenance of SCC.

To ablate USP28 during tumour initiation, we used CRISPR/Cas9 mediated gene targeting. To induce primary lesions in the lungs of mice, we used a constitutive *Rosa26^Sor^-CAGG-Cas9-IRES-GFP* transgenic mouse strain and intratracheally infected these mice at 8 weeks of age with adeno-associated virus (AAV) virions containing sgRNA targeting sequences to inactivate Tp53 (*p53*^Δ^), Stk11/Lkb1 *(Lkb1*^Δ^) and introduce the oncogenic mutation G12D, via a repair template, into the KRas locus. We refer to these mice as KP (*Kras^G12D^:Tp53*^Δ^) or KPL (*Kras^G12D^; Tp53*^Δ^*; Lkb1*^Δ^). At 12 weeks post intratracheal intubation, KP mice developed ADC tumours as determined by the expression of the ADC marker TTF1 and the absence of the SCC marker KRT5 (Figure S5A and B). Co-depletion of the tumour suppressor *STK11/Lkb1*, in combination with *Tp53* and *KRas* targeting, resulted in the development of both major NSCLC entities, ADC (TTF1+/ΔNp63-/KRT5-) and SCC (TTF-1-/ΔNp63+/KRT5+) (Figure 5C and D and Figure S5C). Loss of *Stk11/Lkb1* in KPL mice dramatically shortened overall survival compared to that of KP mice (Figure S5D). Evaluation of USP28 abundance, estimated by IHC, demonstrated an increase in USP28 protein in SCC tumours compared to ADC tumours within the same KPL animal (Figure 5C).

**Figure 5:**
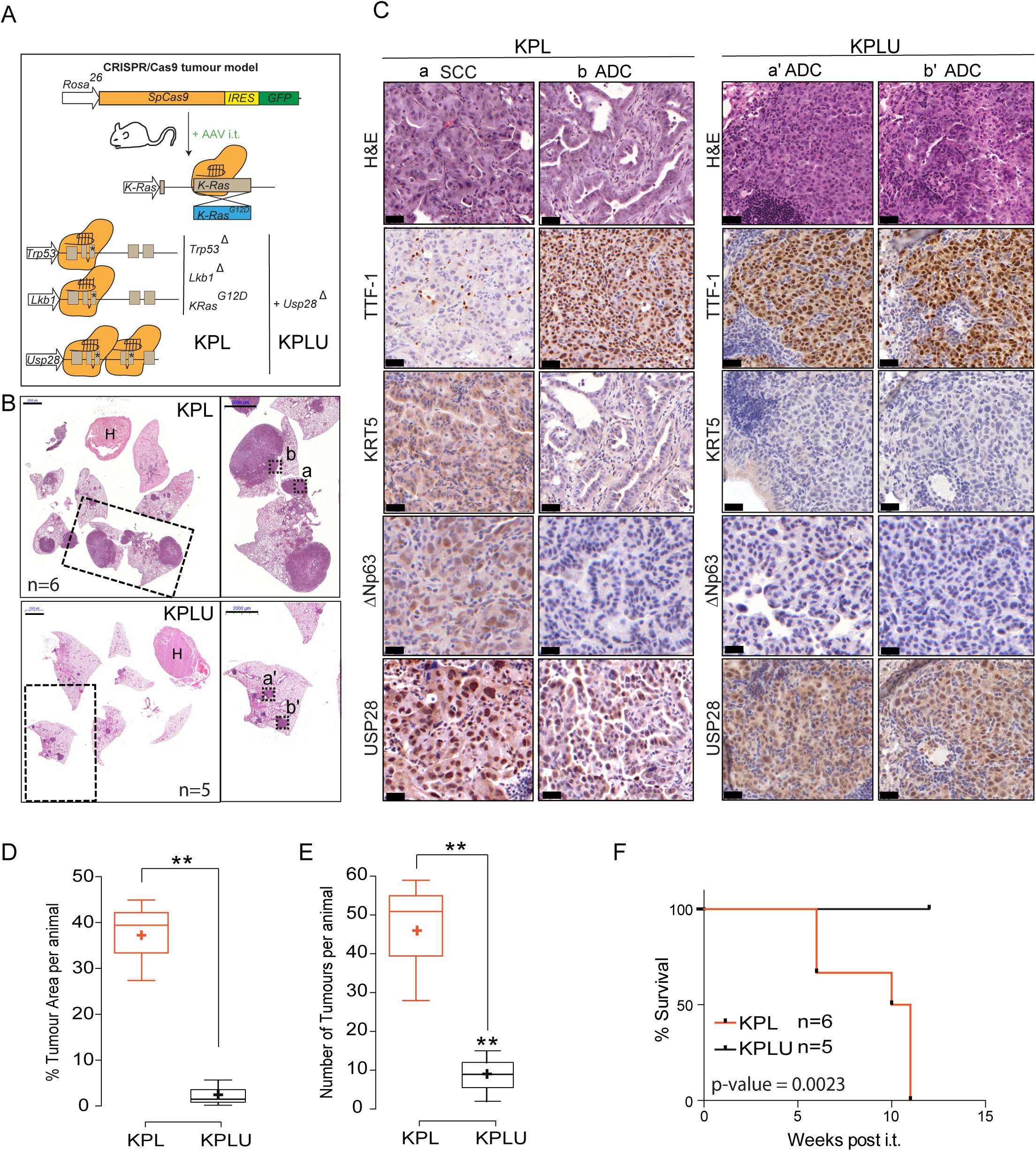
Usp28 is required for SCC tumour formation *in vivo* and affects overall tumour burden. A. Schematic diagram of CRISPR/Cas9 mediated tumour modelling and targeting of *p53*^Δ^*; Lkb1*^Δ^*; KRas^G12D^*(KPL) or *Usp28*^Δ^*; p53*^Δ^*; Lkb1*^Δ^*:KRas^G12D^*(KPLU) mouse lines in *Rosa26Sor-CAGG-Cas9-IRES-GFP* mice, B. Representative haematoxylin and eosin (H&E) staining of tumour bearing animals 12 weeks post intratracheal infection. Boxes indicate highlighted tumour areas in C) (a, b, a’ and b’). Scale bar = 2000µm; nKPL= 6 and nKPLU = 5 C. Representative IHC staining for ADC (TTF-1) and SCC (KRT5 and ΔNp63) marker expression as well as Usp28 abundance in KPL (n=6) and KPLU (n=5) lung tumours. Scale bar: 20µm D. Quantification of % tumour area (normalised to total lung area) in KPL (n=6) and KPLU (n=5) animals. E. Quantification of lung tumour numbers in KPL (n=6) and KPLU (n=6) animals. F. Kaplan-Meier survival curves comparing KPL (n=6) and KPLU (n=5) animals upon AAV intratracheal infection. The p-value was calculated using a log-rank test. In box plots, the centre line reflects the median, the cross represents the mean and the upper and lower box limits indicates the first and third quartile. Whiskers extend 1.5x the IQR. p-values were calculated using two-tailed t-test. **p-value < 0.01. see also Figure S5.

To test the role of USP28 in tumour induction, we included two sgRNA targeting USP28 into the experimental cohort of KLP mice (*USP28*^Δ,^ referred to as KPLU). Concomitant targeting of USP28 at the time of tumour induction significantly affected NSCLC formation (Figure 5B and C). Both, total tumour area as well as tumour number per animal, were significantly reduced in KPLU compared to KPL mice (Figure 5E and F). In stark contrast to KPL mice, tumours occurring in KPLU mice exclusively expressed ADC markers, while no SCC tumours were observed. Consistent with this effect, the survival of KPLU mice was significantly prolonged compared to KPL mice (Figure 5D). Tumours developing in KPL mice showed expression of USP28 and ΔNp63, while USP28 was strongly reduced in isolated tumours from KPLU mice, and ΔNp63 was not detectable (Figure S5F).

Taken together, the data show that USP28 is required for induction of lung SCC.

### ΔNp63-driven SCC cells of various tissues are vulnerable to USP28 depletion

Chromosomal amplification and increased gene expression of ΔNp63 is very common in lung SCC (Figure 7A) when compared to cervix, head-and-neck, oesophagus or pancreatic tumours (Figure 7A). Nevertheless, ΔNp63 expression is increased in SCC of irrespective of ‘tissue of origin’. Therefore, we were wondering if USP28 is also upregulated in these SCC, too. Publicly available expression data sets revealed a significant upregulation of *TP63* and *USP28* in cancer samples from cervix, oesophagus, head-and-neck or lung SCC compared to non-transformed samples (Figure 1 and 7B and C). Patients with an increased expression of *USP28* showed a significantly shortened overall survival in cervix and head-and-neck tumours, while expression of Δ*Np63* significantly shortened life expectancy in pancreatic cancer (Figure S7A and B).

**Figure 7:**
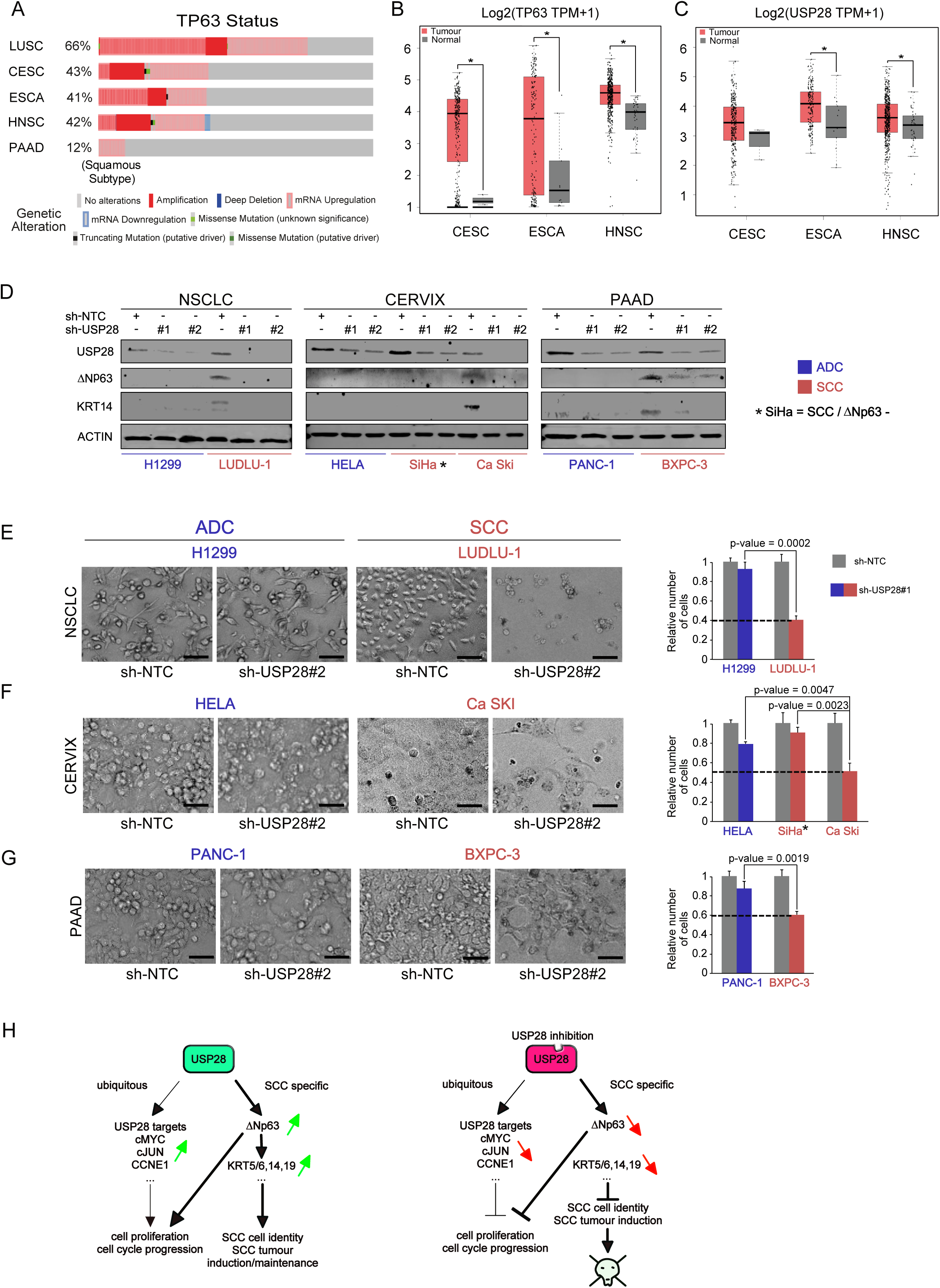
ΔNp63-driven SCC cells of various tissues are vulnerable to USP28 depletion. A. Analysis of occurring TP63 genetic alterations in lung squamous (LUSC), cervical (CESC), oesophagus (ESCA), head and neck (HNSC) and pancreatic (PAAD) tumours. B. Expression of TP63 (left) in human CESC (n=306), ESCA (n=182) and HNSC (n=519) and normal non-transformed tissue (normal CESC=3, normal ESCA=13 and normal HNSC=44). p-values were calculated with a one-way ANOVA comparing normal tissue tumours. TPM: transcript count per million mapped reads C. Expression of USP28 (left) in human CESC (n=306), ESCA (n=182) and HNSC (n=519) and normal non-transformed tissue (normal CESC=3, normal ESCA=13 and normal HNSC=44). p-values were calculated with a one-way ANOVA comparing normal tissue tumours. TPM: transcript count per million mapped reads D. Immunoblot of control (sh-NTC) and two independent shRNA targeting USP28 (sh-USP28#1 and #2) for ΔNp63, KRT14 and USP28 protein abundance in H1299, LUDLU-1, HELA, SiHa, Ca Ski, PANC-1 and BXPC-3 (ACTIN as loading control), E. Cells were seeded at equal cell density and counted after five days, Brigth field images of control or sh-USP28#2 infected H1299 and LUDLU-1 cells before quantification. Relative number of sh-USP28#2 cells compared with the sh-NTC control cells. p-values were calculated using two-tailed t-test statistical analysis. Scale bar = 30µm F. Cells were seeded at equal cell density and counted after five days, Brigth field images of control or sh-USP28#2 infected HELA and Ca Ski cells before quantification. Relative number of sh-USP28#2 cells compared with the sh-NTC control cells, For the quantification, SiHa (Figure S7F) was included, p-values were calculated using two-tailed t-test statistical analysis. Scale bar = 30µm G. Cells were seeded at equal cell density and counted after five days, Brigth field images of control or sh-USP28#2 infected PANC-1 and BXPC3 cells before quantification. Relative number of sh-USP28#2 cells compared with the sh-NTC control cells. p-values were calculated using two-tailed t-test statistical analysis. Scale bar = 30µm H. Schematic Model All quantitative data are represented as mean ± SD (n=3). p-values were calculated using two-tailed t-test statistical analysis; *p-value < 0.05, **p-value < 0.01, *** p-value < 0.001; see also Figure S7.

To investigate a therapeutic potential of targeting the USP28-ΔNp63 axis in SCC cells of different origins, we used a set of human cancer cell lines, comprising the pancreas lines PANC-1 (ADC) and BXPC-3 (SCC), cervical cancer cell lines HeLa (ADC), SiHa and Ca-Ski (SCC), the head-and-neck cell line Detroit 562 (SCC) and the lung cell lines H1299 (ADC) and LUDLU-1 (SCC). While USP28 was readily detectable by immunoblot in all cell lines (Figure S7C), only the SCC lines Ca-Ski, BXPC3 and the head-and-neck cell line Detroit 562 expressed detectable amounts of endogenous ΔNp63. In the cervical line SiHa, despite being of SCC origin, endogenous ΔNp63 was not detectable (Figure S7C).

Next, we targeted USP28 by two independent shRNA sequences, and analysed the knock-down efficacy by immunoblotting (Figure 7D). Both shRNAs induced a significant decrease in USP28 protein levels (Figure 7D). While SCC cell lines infected with a non-targeting control shRNA expressed ΔNp63 and the downstream target KRT14, knock down of USP28 reduced ΔNp63 protein levels, along with KRT14 (Figure 7C), consistent with observed USP28 effects in A431 cells (Figure S4). SCC are uniquely dependent on the expression of ΔNp63, therefore, we next analysed the effect of USP28 knock down on ADC and SCC cell line proliferation (Figure 7E). Control or USP28-depleted ADC and SCC cells were seeded at equal cell density and counted after five days. Proliferation of the ADC cells HeLa, PANC-1 and H1299 was only weakly affected (Figure 7E). In cells of SCC origin expressing ΔNp63, Ca-Ski, BXPC3, Detroit 562 and LUDLU-1, demonstrated a growth disadvantage upon knock-down of USP28 by shRNA (Figure 7D and S7D and E). An exception was the SCC cell line SiHa, which does not express ΔNp63 (Figure 7D and S7F).

USP28 has several targets, including the transcription factors cMYC, cJUN and NOTCH (Diefenbacher et al., 2015; Diefenbacher et al., 2014; Popov et al., 2007a; Schulein-Volk et al., 2014), but the transcriptional responses in SCC cells are dominated by its effects on ΔNp63, arguing that this axis establishes a unique dependence and hence may open a wide therapeutic window for targeting SCC of various tissue origins via USP28 (Figure 7H).

## Discussion

SCC tumours are among the genetically most complex entities (Cancer Genome Atlas Research, 2012). Driver mutations can vary widely, ranging from activating mutations in members of the MAPK-pathway, the PI3K-pathway or RTKs, to gene amplifications in several loci, including the Δ*Np63* gene (Cancer Genome Atlas Research, 2012). SCC tumours have in common is an inherent dependence on ΔNp63 expression (Bergholz and Xiao, 2012; Rocco et al., 2006). Previous work has established the role of ΔNp63 as a master transcription factor and regulator of SCC identity (Abraham et al., 2018; Hamdan and Johnsen, 2018; Somerville et al., 2018). SCC tumour cells are addicted to ΔNp63 expression (Somerville et al., 2018; Vivanco, 2014), as depletion of ΔNp63 is not tolerated by these tumours and leads to rapid tumour regression (Ramsey et al., 2013). Therefore, targeting ΔNp63, either directly or by altering its protein abundance, appears promising for therapy of SCC tumours.

Most transcription factors, including ΔNp63 (Abraham et al., 2018; Dang et al., 2017; Lambert et al., 2018), are considered as ‘non-druggable’ targets as their structure does not provide suitable domains for small molecule interactions. Modulation of their abundance by targeting mechanisms that control protein stability presents a viable option (Liu et al., 2015; Wang et al., 2018). ΔNp63 is tightly regulated by the ubiquitin-proteasome-system and ubiquitylated by various E3-ligases (Armstrong et al., 2016). In a tumour, this mechanism is frequently non-functional, leading to the accumulation of ΔNp63 protein (Ruiz et al., 2019) and results of the current study).

In this study, we explored the possibility to control the abundance of ΔNp63 protein via the deubiquitylase USP28. We found that USP28 is frequently upregulated in human SCC tumours and is often co-expressed with ΔNp63. In contrast, loss or mutation of USP28 is rare and low mRNA expression coincides with better overall survival of patients. We found that USP28 regulates the abundance of ΔNp63 by direct binding and by catalysing the removal of K48-linked ubiquitin chains from the ΔNp63 protein. Loss of USP28 reduced SCC cell proliferation in a ΔNp63-dependent manner, and interfered with the expression of marker genes that define the SCC lineage to a similar extend as targeting ΔNp63 directly. For example, cytokeratin genes commonly expressed in SCC, such as KRT5, KRT14 or KRT19, were similarly downregulated upon loss of ΔNp63 or USP28. The observed effects in gene expression in the USP28 shRNA cell lines were a direct consequence of reduced ΔNp63 protein abundance as expression of exogenous ΔNp63 was able to restore SCC marker expression and cell proliferation in the USP28 targeted cells. Since depletion of USP28 phenocopied the effect of ΔNp63 ablation, targeting USP28 appeared as a valid surrogate molecule to target ΔNp63 in SCC tumours.

In strong support of this notion, targeting USP28 in a model of lung SCC (“KPL mice”) using the CRISPR/Cas9 system abrogated SCC tumour formation. While KPL mice developed both NSCLC entities, ADC and SCC, loss of USP28 strongly affected overall tumour induction and blocked SCC formation. This indicates that SCC are strictly dependent on USP28 expression, consistent with their dependence on ΔNp63 function.

While the interaction between USP28 and ΔNp63 is independent of the catalytic site within the DUB, our results show that the active cysteine at position 171 is essential to facilitate the stabilising effect on the ΔNp63 protein. Mutation of this cysteine residue strongly reduced the ability of USP28 to deubiquitylate ΔNp63. Therefore, inhibiting the catalytic activity of USP28 is likely to be a suitable mechanism to target ΔNp63 in SCC tumours. That the DUB-family presents a viable therapeutic option was tested by using a pan-DUB inhibitor PR-619. SCC cells exposed to PR-619 showed a strong decrease of ΔNp63 levels. USP25/28 dual specific as well as specific inhibitors for other DUBs have been identified and first-generation inhibitors have been reported (Wrigley et al., 2017). Given the crucial dependency of SCC tumour from different tissues on ΔNp63, including those of head-and-neck, cervix, oesophagus, vulva, skin and lung, targeting the dependence of ΔNp63 on USP28, revealed in this study, is a promising strategy to target this tumour entity independent of tissue origin.

## Supporting information

Consensus Squamous Marker Signature Genes

Supplementary table 1 USP28 and ???Np63 deregulated genes Garcia et al

Supplementary table 3 Materials and Resources Garcia et al

## Acknowledgements

We are grateful to the animal facility and Barbara Bauer at the Biocenter University Würzburg. C.P.G. and O.H. are supported by the German Cancer Aid via grant 70112491, M.E. is supported by the TransOnc priority program of the German Cancer Aid within grant 70112951 (ENABLE), M.R. is funded by the DFG-GRK 2243 and IZKF B335. M.E.D. and A.O. are funded by the German Israeli Foundation grant 1431.

## Author contributions

Conceptualization: C.P.G., M.E., M.E.D.; Methodology: C.P.G. (in vitro) and O.H. (*in vivo*), C.P.G. (Biochemistry), M.R. (TUBE), F.B. (FACS); Formal analysis: S.W. and C.P.G. (Bioinformatics), M.C., M.R. and M.E.D. (Pathology); Investigation: C.P.G., O.H., M.R., T.F., F.B., M.R., M.E.D. Resources: M. C., M.R., M.E.D.; Writing-original draft: M.E.D.; Writing-review and editing: C.P.G., O.H., M.R., A.O., M.E., M.E.D.; Supervision: M.E. and M.E.D.; Funding acquisition: M.E.D.

## Declaration of Interests

The authors declare no competing interests.

## Experimental Model and Subject Details

### Tissue culture, transfection, infection and reagents

Cells were plated on Greiner petri dishes and maintained at 37°C, 95% relative humidity and 5% CO2 for optimal growth conditions. All cell lines were obtained from ATCC or ECACC. A-431, PANC-1, SiHa, Ca SKI, DETROIT 562 and HEK-293T cells were cultured in DMEM (Gibco;) supplemented with 10% fetal bovine serum (FCS) and 1% Pen-Strep. Ludlu-1, NCI-H1299 and BxPC-3 cells were cultured in RPMI 1640 (Gibco) supplemented with 10% FCS, 1% GlutaMAX and 1% Pen Strep.

For DNA transfection, a mix of 2,5μg plasmid DNA, 200μl free medium and 5μl PEI was added into the 6-well dish medium (60% confluence), after 6h incubation at 37°C the medium was changed to full supplemented medium. For DNA infection, AAVs or Lentiviruses (MOI=10) were added to the cell medium in the presence of polybrene (5μg/ml) and incubating at 37°C for 96h. After incubation, infected cells were selected with 2,5 μg/ml Puromycin for 72h, 250µg/ml Neomycin for 2 weeks or FACS-sorted for RFP/GFP positive cells (FACS Canto II BD).

Except when a different concentration was expressly indicated, the next reagents were dissolved in Dimethyl sulfoxide (DMSO) or 70% ethanol and added to the cells at the following concentrations: Cycloheximide (CHX; 100 μg/ml), doxycycline (DOX; 1μg/ml), Tandem ubiquitin binding entity (TUBE; 100 μg/ml).

### RT-PCR

RNA was isolated with Peq GOLD Trifast (Peqlab), following the manufacturer’s instructions. RNA was reverse transcribed into complementary DNA (cDNA) using random hexanucleotide primers and M-MLV reverse transcriptase (Promega). Quantitative RT-PCR was performed using QPCR SYBR Green mix (ABgene) on the instrument ‘Step One Realtime Cycler’(ABgene) RT-PCR was performed using the next Program: 95°C for 15 min., 40x [95°C for 15 sec., 60°C for 20 sec. and 72°C for 15 sec.], 95°C for 15 sec. and 60°C for 60 sec. Relative expression was generally calculated with ΔΔCt relative quantification method using the expression of B-ACTIN as House-keeping gene. For mouse USP28 mRNA expression in A431, relative expression was calculated using ΔCt. For all the experiments, melt curve was performed. Primers used for this publication are listed,

### Immunological Methods

Cells were lysed in RIPA lysis buffer (20 mM Tris-HCl pH 7.5, 150 mM NaCl, 1mM Na2EDTA, 1mM EGTA, 1% NP-40 and 1% sodium doxycholate), containing proteinase inhibitor (1/100) by sonication using Branson Sonifier 250 with a duty cycle at 20%, output control set on level 2 and the timer set to 1 min (10 sonication cycles per sample). Protein concentration was quantified using Bradford assay. After mixing 1 ml of Bradford reagent with 1μl of sample, the photometer was used to normalize the protein amounts with a previously performed bovine serum albumin (BSA) standard curve. The quantified protein (40μg) was boiled in 5x Laemmli buffer (312.5mM Tris-HCl pH 6.8, 500 mM DTT, 0.0001% Bromphenol blue, 10% SDS and 50% Glycerol) for 5 min and separated on 10% Tris-gels in Running buffer (1.25M Tris base, 1.25M glycine and 1% SDS). After separation, protein was transferred to Polyvinylidene difluoride membranes (Immobilon-FL) in Transfer Buffer (25mM Tris base, 192mM glycine and 20% methanol) and then, incubated with blocking buffer (0.1% casein, 0.2xPBS and 0.1% Tween20) for 45 Min at RT. After Blocking, membranes were incubated with indicated Primary Abs (1/1000 dilution in a buffer composed by 0.1% casein, 0.2x PBS and 0.1% Tween20) for 4H at room temperature (RT). Secondary Abs (1/10000 dilution in a buffer composed by 0.1% casein, 0.2x PBS, 0.1% Tween20 and 0.01% SDS) were incubated for 1H at RT. Membranes were recorded with Odyssey® CLx Imaging System or iBright™ FL1000 Imaging System. Analysis and quantifications of protein expression was performed using Image Studio software (Licor Sciences). Data are presented as mean and error bars as standard deviation of the biological replicates. Significance was calculated with Excel using heteroscedastic two-tailed Student’s t-tests. Immunoprecipitation was performed using 0,25 mg of Pierce™ Protein A/G Magnetic Beads (ThermoFisher), 1μg of the specific Ab and 500μg of protein lysate. For endogenous Co-Immunoprecipitations, beads were incubated with Rabbit IgG as a control for specificity. For TUBE assay, 100 μg/ml of recombinant expressed GST-4x UIM-Ubiquilin fusion protein was mixed with the cell lysate before protein extraction and then processed for Western Blot detection. Antibodies used for this publication are listed.

### Immunofluorescence

Standard procedures were used for immunohistochemistry (IHC) and immunofluorescence (IF). For antibodies used in IHC or IF, manufacturer’s manuals and instructions were used regarding concentration or buffer solutions. All primary antibodies were incubated over night at 4°C, followed by washing thrice with PBS, and subsequent incubation with the secondary antibody for 1 hour at room temperature. IHC slides were imaged using Pannoramic DESK scanner and analyzed using Case Viewer software (3DHISTECH). For samples stained with IF, tissue-samples/cells were counterstained with 5 μg/ml DAPI, to highlight nuclei, for 15 minutes after secondary antibody application. Stained samples were mounted with Mowiol®40-88 and imaged using a FSX100 microscopy system (Olympus).

### FACS, growth curves and colony formation assays

FACS analysis and quantification was performed with FACS Canto II (BD) flow cytometer and results were quantified and visualized using FlowJo software. For cell cycle analysis, cells were fixed using 70% ethanol and stained with Propidium Iodide (Sigma Aldrich) at 4°C for 30 Minutes. For cell number quantification, Cell lines were seeded at 5-10% confluence in 24-well dishes. After 3.5, 5 or 7 days, tumour cells were trypsinized and counted with Invitrogen Countess II FL. Colony assays were performed staining 70% ethanol fixed cells with 0,5%. All the Graphics except box-plots were generated using Excel (Microsoft). Data was presented as mean and error bars as standard deviation of 3 independent biological replicates.

### Site-Directed Mutagenesis sgRNA and shRNA Design

sgRNAs were designed using the CRISPRtool (https://zlab.bio/guide-design-resources). shRNA sequences were designed using SPLASH-algorithm (http://splashrna.mskcc.org/) (Pelossof et al., 2017) or the RNAi Consortium/Broad Institute (www.broadinstitute.org/rnai-consortium/rnai-consortium-shrna-library). For site directed mutagenesis, guidance and instructions from the kit GeneEditor™ in vitro Site-Directed Mutagenesis System (Promega) were followed

### AAV and lentivirus production and purification

Virus was packaged and synthetized in HEK293-T cells seeded in 15cm-dishes. For AAV production, cells (70% confluence) were transfected with the plasmid of interest (10μg), pHelper (15μg) and pAAV-DJ or pAAV-2/8 (10μg) using PEI (70μg). After 96h, the cells and medium of 3 dishes were transferred to a 50 ml Falcon tube together with 5 ml chloroform. Then, the mixture was shaken at 37 C for 60 min and NaCl (1M) was added to the mixture. After NaCl is dissolved, the tubes were centrifuged at 20,000 x g at 4°C for 15 min and the chloroform layer was transferred to another Falcon tube together with 10% PEG8000. As soon as the PEG800 is dissolved, the mixture was incubated at 4C overnight and pelleted at 20,000 x g at 4°C for 15 min. The pellet was resuspended in PBS with MgCl2 and 0.001% pluronic F68, then, the virus was purified using 1X Chloroform and stored at −80C. AAV viruses were titrated using Coomassie Staining and RT-PCR using AAV-ITR sequence specific primers.

For Lentivirus production, HEK293-T cells (70% confluence) were transfected with the plasmid of interest (15μg), pPAX (10μg) and pPMD2 (10μg) using PEI (70μg). After 96h, the medium containing lentivirus was filtered (0.45uM) and stored at −80C.

### In Vivo Experiments and Histology

All *in vivo* experiments were approved by the Regierung Unterfranken and the ethics committee under the license numbers 2532-2-362, 2532-2-367 and 2532-2-374. The mouse strains used for this publication are listed.

Adult mice (around eight Weeks old) were anesthetized with Isoflurane and intratracheally intubated with 50 μl AAV virus (3 × 10^7^ PFU) diluted in PBS. Viruses were quantified using Coomassie staining protocol (Kohlbrenner et al., 2012) and cells were quantified with Invitrogen Countess II FL. For Intratracheal instillation, a gauge 24 catheter was introduced to the trachea and cells or virus previously isolated were pipetted to the top of the catheter. During animal breathing, cells and virus were delivered into the lungs. At the indicated time point, animals were sacrificed by cervical dislocation and lungs were dissected and later fixed using 5% NBF. Samples were embedded in paraffin and sectioned at 6 μm using the microtome (Leica). Before staining, slides were de-paraffinized and rehydrated using the next protocol: 2x 5 min. in Xylene, 2x 3 min. in EtOH (100 %), 2x 3 min. in EtOH (95 %), 2x 3 min. in EtOH (70%), 3 min. in EtOH (50 %) and 3 min. in H_2_O. Slides were stained with haematoxylin and eosin or Immunohistochemistry (IHC) using the reported antibodies. For all staining variants, slides were mounted with 200μl of Mowiol® 40-88 covered up by a coverslip on top. IHC slides were recorded using Pannoramic DESK scanner and analysed using Case Viewer software (3DHISTECH). IF samples were recorded using FSX100 microscopy system (Olympus). Quantification of the tumour area and number of tumours were performed using ImageJ as previously described(Jensen, 2013).

### Human lung cancer samples

Lung cancer samples were obtained, stored, and managed by Pathology Department Cordoba, Pathology Department University Hospital Würzburg and U.S. Biomaxx (Lung microarray slides; slide LC2083). Informed consent was obtained for each human sample. Human tissue samples were stained against USP28 and ΔNP63. For quantification purposes, staining intensity was graded from 0 (no staining) up to 3 (staining intensity >66%) by three independent pathologists. Box-plots were generated using BoxPlotR online tool (http://shiny.chemgrid.org/boxplotr/). The centre line reflects the median, the cross represents the mean and the upper and lower box limits indicates the first and third quartile, respectively. Whiskers extend 1.5x the IQR and outliers are marked as dots. The significance was calculated using t-test.

### RNA-sequencing

RNA sequencing was performed with Illumina NextSeq 500 as described previously (Buchel et al., 2017).RNA was isolated using ReliaPrep™ RNA Cell Miniprep System Promega kit, following the manufacturer’s instruction manual. mRNA was purified with NEBNext® Poly(A) mRNA Magnetic Isolation Module (NEB) and the library was generated using the NEBNext® UltraTM RNA Library Prep Kit for Illumina, following the manufacturer’s instructions. For the size-selection of the libraries, Agencourt AMPure XP Beads (Beckman Coulter) were used. Library quantification and size determination was performed using Fragment Analyzer (Agilent formerly Advanced Analytical).

## QUANTIFICATION AND STATISTICAL ANALYSIS

### Analysis of publicly available data

All publicly available data and software used for this publication are listed in the key resource table. Oncoprints were generated using cBioportal (Cerami et al., 2012; Gao et al., 2013). Briefly, Oncoprints generates graphical representations of genomic alterations, somatic mutations, copy number alterations and mRNA expression changes. The following studies were used for the analysis: lung adenocarcinoma (“LUAD-TCGA, Provisional”), lung squamous cell carcinoma (“LUSC-TCGA, Provisional”), small cell lung cancer (“U Cologne, Nature 2015”), cervical cancer (“CESC-TCGA, Provisional”), Esophagus cancer (“ESCA-TCGA, Provisional”), head and neck tumours (“HNSC-TCGA, Provisional”) and pancreatic cancer (“PAAD-TCGA, Provisional”)The mRNA expression Z-score threshold was 1.5.

Box plots using TCGA and GTEx data were generated using the online tool GEPIA (Tang et al., 2017). In box plots, the central line shows the second quartile and the upper and lower border of the box reflect the 25th and 75th percentiles, respectively. Whiskers represent 1.5x of the interquartile range (IQR) and individual data points are represented by black dots. The differential analysis was based on: “TCGA tumours vs (TCGA normal + GTEx normal)”, whereas the expression data were log_2_(TPM+1) transformed and the log_2_FC was defined as median(tumour) – median(normal). p-values were calculated with a one-way ANOVA comparing tumour with normal tissue.

Correlation USP28 and TP63 expression in tumour and normal tissue was calculated using GEPIA’s software, The analysis was based on the expression of the following datasets: “TCGA tumours”, “TCGA normal” and “GTEx normal”. The expression of USP28, TP63 and the gene set “Squamous Cancer Markers” (consensus list of up-regulated genes for squamous tumours based on previous publications (Ferone et al., 2016; Mukhopadhyay et al., 2014; Wilkerson et al., 2010; Xu et al., 2014)) were used for the calculation of the Spearman’s correlation coefficents and significance by GEPIA’s software

The comparison of gene expression (using the data-set Squamous cancer markers) across multiple tumour entities was done using GEPIA software based on the dataset “TCGA tumours”. The color code reflects the median expression of a gene in a tumour type, normalised with the maximum median expression across all different tumour types (row-wise Z-score).

Genomic signature comparing primary human lung tumour was performed using UCSC Xena (https://doi.org/10.1101/326470) based on the dataset “TCGA tumours”.

Kaplan-Meier curves were estimated with the KM-plotter (Lanczky A. et al 2016), cBioportal (Gao et al. Sci. Signal. 2013 & Cerami et al. Cancer Discov. 2012) and R2: Genomics Analysis and Visualization Platform (http://r2.amc.nl). The KM-plotter was used to analyse overall survival of lung cancer patients (Figure 1) based on gene expression data from microarrays obtained from GEO, caBIG and TCGA (REF). For R2: Genomics Analysis and Visualization Platform (http://r2.amc.nl), overall survival and gene expression data were obtained from TCGA (Figure S7). For the survival analysis of USP28 altered samples (mutation or deep deletion), cBioportal was used to calulate disease-free survival using the dataset “Lung Squamous Cell Carcinoma (LUSC-TCGA, Provisional)” (Figure S1). p-values were computed using a log-rank test.

### RNA-sequencing analysis

Fastq files were generated using Illuminas base calling software GenerateFASTQ v1.1.0.64 and overall sequencing quality was analyzed using the FastQC script. Reads were aligned to the human genome (hg19) using Tophat v2.1.1 (Kim et al., 2013) and Bowtie2 v2.3.2 (Langmead and Salzberg, 2012) and samples were normalised to the number of mapped reads in the smallest sample. For differential gene expression analysis, reads per gene (Ensembl gene database) were counted with the “summarizeOverlaps” function from the R package “GenomicAlignments” using the “union”-mode and non-or weakly expressed genes were removed (mean read count over all samples <1). Differentially expressed genes were called using edgeR (Robinson et al., 2010) and resulting p-values were corrected for multiple testing by false discovery rate (FDR) calculations.

### GSEA

For gene set enrichment analyses (GSEA, (Mootha et al., 2003; Subramanian et al., 2005), five gene sets were generated: “Genes Down-regulated sh-USP28” (q-value<0.05, log_2_FC<-1.5), “Gene Up-regulated sh-USP28” (q-value<0.05, log_2_FC>1.5), “Genes Down-regulated sh-ΔNP63” (q-value<0.05, log_2_FC<-3),“Genes Up-regulated sh-ΔNP63” (q-value<0.05, log_2_FC>3) and “Squamous Cancer marker” (consensus list of up-regulated genes for squamous tumors). Genes included in each gene set are reported in Supp. Table 1 and 2. GSEA analyses were done with signal2Noise metric and 1000 permutations.

### Gene ontology analysis

Gene ontology analysis were performed with PANTHER (Mi et al., 2019) using the “Statistical overrepresentation test” tool with default settings. Three gene sets were generated based on the RNA-seq data and analysed: “Genes Down-regulated sh-USP28” (q-value<0.05, log_2_FC>1.5), “Genes Down-regulated sh-ΔNP63” (q-value<0.05, log_2_FC<-1.5).

### Venn diagrams and pair-wise correlations

Venn diagrams were visualised using the online tool: http://bioinformatics.psb.ugent.be/webtools/Venn/. p-values were calculated with a hypergeometric test using the online tool at http://nemates.org. p-values for correlation coefficents were calculated using two-tailed Student’s t-tests.

## DATA AND SOFTWARE AVAILABILITY

RNA-sequencing data is available at the Gene Expression.

## Contact for reagent and resource sharing

Further information and requests for resources and reagents should be directed to and will be fulfilled by the Lead Contact, Markus E. Diefenbacher.

## Notes

**Financial Support:** C.P.G. and O.H. are supported by the German Cancer Aid via grant 70112491. M.E. is supported by the TransOnc priority program of the German Cancer Aid within grant 70112951 (ENABLE). M.R. is funded by the DFG-GRK 2243 and IZKF B335. M.E.D. and A.O. are funded by the German Israeli Foundation grant 1431.

**Conflict of Interest:** The authors declare no potential conflicts of interest.

## References

Abraham, C.G., Ludwig, M.P., Andrysik, Z., Pandey, A., Joshi, M., Galbraith, M.D., Sullivan, K.D., and Espinosa, J.M. (2018). DeltaNp63alpha Suppresses TGFB2 Expression and RHOA Activity to Drive Cell Proliferation in Squamous Cell Carcinomas. Cell Rep 24, 3224–3236.

Antonini, D., Rossi, B., Han, R., Minichiello, A., Di Palma, T., Corrado, M., Banfi, S., Zannini, M., Brissette, J.L., and Missero, C. (2006). An autoregulatory loop directs the tissue-specific expression of p63 through a long-range evolutionarily conserved enhancer. Mol Cell Biol 26, 3308–3318.

Armstrong, S.R., Wu, H., Wang, B., Abuetabh, Y., Sergi, C., and Leng, R.P. (2016). The Regulation of Tumor Suppressor p63 by the Ubiquitin-Proteasome System. Int J Mol Sci 17.

Bergholz, J., and Xiao, Z.X. (2012). Role of p63 in Development, Tumorigenesis and Cancer Progression. Cancer Microenviron 5, 311–322.

Buchel, G., Carstensen, A., Mak, K.Y., Roeschert, I., Leen, E., Sumara, O., Hofstetter, J., Herold, S., Kalb, J., Baluapuri, A., et al. (2017). Association with Aurora-A Controls N-MYC-Dependent Promoter Escape and Pause Release of RNA Polymerase II during the Cell Cycle. Cell Rep 21, 3483–3497.

Cancer Genome Atlas Research, N. (2012). Comprehensive genomic characterization of squamous cell lung cancers. Nature 489, 519–525.

Cerami, E., Gao, J., Dogrusoz, U., Gross, B.E., Sumer, S.O., Aksoy, B.A., Jacobsen, A., Byrne, C.J., Heuer, M.L., Larsson, E., et al. (2012). The cBio cancer genomics portal: an open platform for exploring multidimensional cancer genomics data. Cancer Discov 2, 401–404.

Craig, A.L., Holcakova, J., Finlan, L.E., Nekulova, M., Hrstka, R., Gueven, N., DiRenzo, J., Smith, G., Hupp, T.R., and Vojtesek, B. (2010). DeltaNp63 transcriptionally regulates ATM to control p53 Serine-15 phosphorylation. Molecular cancer 9, 195.

Cremona, C.A., Sancho, R., Diefenbacher, M.E., and Behrens, A. (2016). Fbw7 and its counteracting forces in stem cells and cancer: Oncoproteins in the balance. Semin Cancer Biol 36, 52–61.

Dang, C.V., Reddy, E.P., Shokat, K.M., and Soucek, L. (2017). Drugging the ‘undruggable’ cancer targets. Nat Rev Cancer 17, 502–508.

de Bie, P., and Ciechanover, A. (2011). Ubiquitination of E3 ligases: self-regulation of the ubiquitin system via proteolytic and non-proteolytic mechanisms. Cell Death Differ 18, 1393–1402.

Deyoung, M.P., and Ellisen, L.W. (2007). p63 and p73 in human cancer: defining the network. Oncogene 26, 5169–5183.

Diefenbacher, M.E., Chakraborty, A., Blake, S.M., Mitter, R., Popov, N., Eilers, M., and Behrens, A. (2015). Usp28 counteracts Fbw7 in intestinal homeostasis and cancer. Cancer Res 75, 1181–1186.

Diefenbacher, M.E., Popov, N., Blake, S.M., Schulein-Volk, C., Nye, E., Spencer-Dene, B., Jaenicke, L.A., Eilers, M., and Behrens, A. (2014). The deubiquitinase USP28 controls intestinal homeostasis and promotes colorectal cancer. J Clin Invest 124, 3407–3418.

Ferone, G., Song, J.Y., Sutherland, K.D., Bhaskaran, R., Monkhorst, K., Lambooij, J.P., Proost, N., Gargiulo, G., and Berns, A. (2016). SOX2 Is the Determining Oncogenic Switch in Promoting Lung Squamous Cell Carcinoma from Different Cells of Origin. Cancer Cell 30, 519–532.

Galli, F., Rossi, M., D’Alessandra, Y., De Simone, M., Lopardo, T., Haupt, Y., Alsheich-Bartok, O., Anzi, S., Shaulian, E., Calabro, V., et al. (2010). MDM2 and Fbw7 cooperate to induce p63 protein degradation following DNA damage and cell differentiation. J Cell Sci 123, 2423–2433.

Gao, J., Aksoy, B.A., Dogrusoz, U., Dresdner, G., Gross, B., Sumer, S.O., Sun, Y., Jacobsen, A., Sinha, R., Larsson, E., et al. (2013). Integrative analysis of complex cancer genomics and clinical profiles using the cBioPortal. Sci Signal 6, pl1.

Gersch, M., Wagstaff, J.L., Toms, A.V., Graves, B., Freund, S.M.V., and Komander, D. (2019). Distinct USP25 and USP28 Oligomerization States Regulate Deubiquitinating Activity. Mol Cell.

Grice, G.L., and Nathan, J.A. (2016). The recognition of ubiquitinated proteins by the proteasome. Cell Mol Life Sci 73, 3497–3506.

Hamdan, F.H., and Johnsen, S.A. (2018). DeltaNp63-dependent super enhancers define molecular identity in pancreatic cancer by an interconnected transcription factor network. Proc Natl Acad Sci U S A 115, E12343–E12352.

Hibi, K., Trink, B., Patturajan, M., Westra, W.H., Caballero, O.L., Hill, D.E., Ratovitski, E.A., Jen, J., and Sidransky, D. (2000). AIS is an oncogene amplified in squamous cell carcinoma. Proceedings of the National Academy of Sciences of the United States of America 97, 5462–5467.

Hjerpe, R., Aillet, F., Lopitz-Otsoa, F., Lang, V., England, P., and Rodriguez, M.S. (2009). Efficient protection and isolation of ubiquitylated proteins using tandem ubiquitin-binding entities. EMBO Rep 10, 1250–1258.

Inamura, K. (2017). Lung Cancer: Understanding Its Molecular Pathology and the 2015 WHO Classification. Front Oncol 7, 193.

Jensen, E.C. (2013). Quantitative analysis of histological staining and fluorescence using ImageJ. Anat Rec (Hoboken) 296, 378–381.

Kohlbrenner, E., Henckaerts, E., Rapti, K., Gordon, R.E., Linden, R.M., Hajjar, R.J., and Weber, T. (2012). Quantification of AAV particle titers by infrared fluorescence scanning of coomassie-stained sodium dodecyl sulfate-polyacrylamide gels. Hum Gene Ther Methods 23, 198–203.

Koster, M.I., Dai, D., Marinari, B., Sano, Y., Costanzo, A., Karin, M., and Roop, D.R. (2007). p63 induces key target genes required for epidermal morphogenesis. Proc Natl Acad Sci U S A 104, 3255–3260.

Lambert, M., Jambon, S., Depauw, S., and David-Cordonnier, M.H. (2018). Targeting Transcription Factors for Cancer Treatment. Molecules 23.

Lau, C.P., Ng, P.K., Li, M.S., Tsui, S.K., Huang, L., and Kumta, S.M. (2013). p63 regulates cell proliferation and cell cycle progressionassociated genes in stromal cells of giant cell tumor of the bone. Int J Oncol 42, 437–443.

Liu, J., Shaik, S., Dai, X., Wu, Q., Zhou, X., Wang, Z., and Wei, W. (2015). Targeting the ubiquitin pathway for cancer treatment. Biochim Biophys Acta 1855, 50–60.

McDade, S.S., Patel, D., and McCance, D.J. (2011). p63 maintains keratinocyte proliferative capacity through regulation of Skp2-p130 levels. J Cell Sci 124, 1635–1643.

Mukhopadhyay, A., Berrett, K.C., Kc, U., Clair, P.M., Pop, S.M., Carr, S.R., Witt, B.L., and Oliver, T.G. (2014). Sox2 cooperates with Lkb1 loss in a mouse model of squamous cell lung cancer. Cell Rep 8, 40–49.

Pelossof, R., Fairchild, L., Huang, C.H., Widmer, C., Sreedharan, V.T., Sinha, N., Lai, D.Y., Guan, Y., Premsrirut, P.K., Tschaharganeh, D.F., et al. (2017). Prediction of potent shRNAs with a sequential classification algorithm. Nat Biotechnol 35, 350–353.

Peschiaroli, A., Scialpi, F., Bernassola, F., El Sherbini el, S., and Melino, G. (2010). The E3 ubiquitin ligase WWP1 regulates DeltaNp63-dependent transcription through Lys63 linkages. Biochem Biophys Res Commun 402, 425–430.

Popov, N., Herold, S., Llamazares, M., Schulein, C., and Eilers, M. (2007a). Fbw7 and Usp28 regulate myc protein stability in response to DNA damage. Cell Cycle 6, 2327–2331.

Popov, N., Wanzel, M., Madiredjo, M., Zhang, D., Beijersbergen, R., Bernards, R., Moll, R., Elledge, S.J., and Eilers, M. (2007b). The ubiquitin-specific protease USP28 is required for MYC stability. Nat Cell Biol 9, 765–774.

Ramsey, M.R., Wilson, C., Ory, B., Rothenberg, S.M., Faquin, W., Mills, A.A., and Ellisen, L.W. (2013). FGFR2 signaling underlies p63 oncogenic function in squamous cell carcinoma. J Clin Invest 123, 3525–3538.

Rocco, J.W., Leong, C.O., Kuperwasser, N., DeYoung, M.P., and Ellisen, L.W. (2006). p63 mediates survival in squamous cell carcinoma by suppression of p73-dependent apoptosis. Cancer Cell 9, 45–56.

Ruiz, E.J., Diefenbacher, M.E., Nelson, J.K., Sancho, R., Pucci, F., Chakraborty, A., Moreno, P., Annibaldi, A., Liccardi, G., Encheva, V., et al. (2019). LUBAC determines chemotherapy resistance in squamous cell lung cancer. J Exp Med 216, 450–465.

Saladi, S.V., Ross, K., Karaayvaz, M., Tata, P.R., Mou, H., Rajagopal, J., Ramaswamy, S., and Ellisen, L.W. (2017). ACTL6A Is Co-Amplified with p63 in Squamous Cell Carcinoma to Drive YAP Activation, Regenerative Proliferation, and Poor Prognosis. Cancer Cell 31, 35–49.

Sauer, F., Klemm, T., Kollampally, R.B., Tessmer, I., Nair, R.K., Popov, N., and Kisker, C. (2019). Differential Oligomerization of the Deubiquitinases USP25 and USP28 Regulates Their Activities. Mol Cell.

Schulein-Volk, C., Wolf, E., Zhu, J., Xu, W., Taranets, L., Hellmann, A., Janicke, L.A., Diefenbacher, M.E., Behrens, A., Eilers, M., et al. (2014). Dual regulation of Fbw7 function and oncogenic transformation by Usp28. Cell Rep 9, 1099–1109.

Somerville, T.D.D., Xu, Y., Miyabayashi, K., Tiriac, H., Cleary, C.R., Maia-Silva, D., Milazzo, J.P., Tuveson, D.A., and Vakoc, C.R. (2018). TP63-Mediated Enhancer Reprogramming Drives the Squamous Subtype of Pancreatic Ductal Adenocarcinoma. Cell Rep 25, 1741–1755 e1747.

Su, X., Chakravarti, D., and Flores, E.R. (2013). p63 steps into the limelight: crucial roles in the suppression of tumorigenesis and metastasis. Nature reviews Cancer 13, 136–143.

Tang, Z., Li, C., Kang, B., Gao, G., Li, C., and Zhang, Z. (2017). GEPIA: a web server for cancer and normal gene expression profiling and interactive analyses. Nucleic Acids Res 45, W98–W102.

Tonon, G., Wong, K.K., Maulik, G., Brennan, C., Feng, B., Zhang, Y., Khatry, D.B., Protopopov, A., You, M.J., Aguirre, A.J., et al. (2005). High-resolution genomic profiles of human lung cancer. Proceedings of the National Academy of Sciences of the United States of America 102, 9625–9630.

Vivanco, I. (2014). Targeting molecular addictions in cancer. Br J Cancer 111, 2033–2038.

Wang, F., Wang, L., Wu, J., Sokirniy, I., Nguyen, P., Bregnard, T., Weinstock, J., Mattern, M., Bezsonova, I., Hancock, W.W., et al. (2017). Active site-targeted covalent irreversible inhibitors of USP7 impair the functions of Foxp3+ T-regulatory cells by promoting ubiquitination of Tip60. PLoS One 12, e0189744.

Wang, L., Xia, W., Chen, H., and Xiao, Z.X. (2019). DeltaNp63alpha modulates phosphorylation of p38 MAP kinase in regulation of cell cycle progression and cell growth. Biochem Biophys Res Commun 509, 784–789.

Wang, X., Liu, Z., Zhang, L., Yang, Z., Chen, X., Luo, J., Zhou, Z., Mei, X., Yu, X., Shao, Z., et al. (2018). Targeting deubiquitinase USP28 for cancer therapy. Cell Death Dis 9, 186.

Westfall, M.D., Mays, D.J., Sniezek, J.C., and Pietenpol, J.A. (2003). The Delta Np63 alpha phosphoprotein binds the p21 and 14-3-3 sigma promoters in vivo and has transcriptional repressor activity that is reduced by Hay-Wells syndrome-derived mutations. Molecular and cellular biology 23, 2264–2276.

Wilkerson, M.D., Yin, X., Hoadley, K.A., Liu, Y., Hayward, M.C., Cabanski, C.R., Muldrew, K., Miller, C.R., Randell, S.H., Socinski, M.A., et al. (2010). Lung squamous cell carcinoma mRNA expression subtypes are reproducible, clinically important, and correspond to normal cell types. Clin Cancer Res 16, 4864–4875.

Winters, I.P., Chiou, S.H., Paulk, N.K., McFarland, C.D., Lalgudi, P.V., Ma, R.K., Lisowski, L., Connolly, A.J., Petrov, D.A., Kay, M.A., et al. (2017). Multiplexed in vivo homology-directed repair and tumor barcoding enables parallel quantification of Kras variant oncogenicity. Nat Commun 8, 2053.

Wrigley, J.D., Gavory, G., Simpson, I., Preston, M., Plant, H., Bradley, J., Goeppert, A.U., Rozycka, E., Davies, G., Walsh, J., et al. (2017). Identification and Characterization of Dual Inhibitors of the USP25/28 Deubiquitinating Enzyme Subfamily. ACS Chem Biol 12, 3113–3125.

Xu, C., Fillmore, C.M., Koyama, S., Wu, H., Zhao, Y., Chen, Z., Herter-Sprie, G.S., Akbay, E.A., Tchaicha, J.H., Altabef, A., et al. (2014). Loss of Lkb1 and Pten leads to lung squamous cell carcinoma with elevated PD-L1 expression. Cancer Cell 25, 590–604.

Yeh, C.H., Bellon, M., Pancewicz-Wojtkiewicz, J., and Nicot, C. (2016). Oncogenic mutations in the FBXW7 gene of adult T-cell leukemia patients. Proc Natl Acad Sci U S A 113, 6731–6736.

Mary Goldman, Brian Craft, Mim Hastie, Kristupas Repečka, Akhil Kamath, Fran McDade, Dave Rogers, Angela N Brooks, Jingchun Zhu, David Haussler (2019). bioRxiv 326470; doi: https://doi.org/10.1101/326470

