## Supplementary Figures 1-7 for "The USP28-ΔNp63 axis is a vulnerability of squamous tumours"

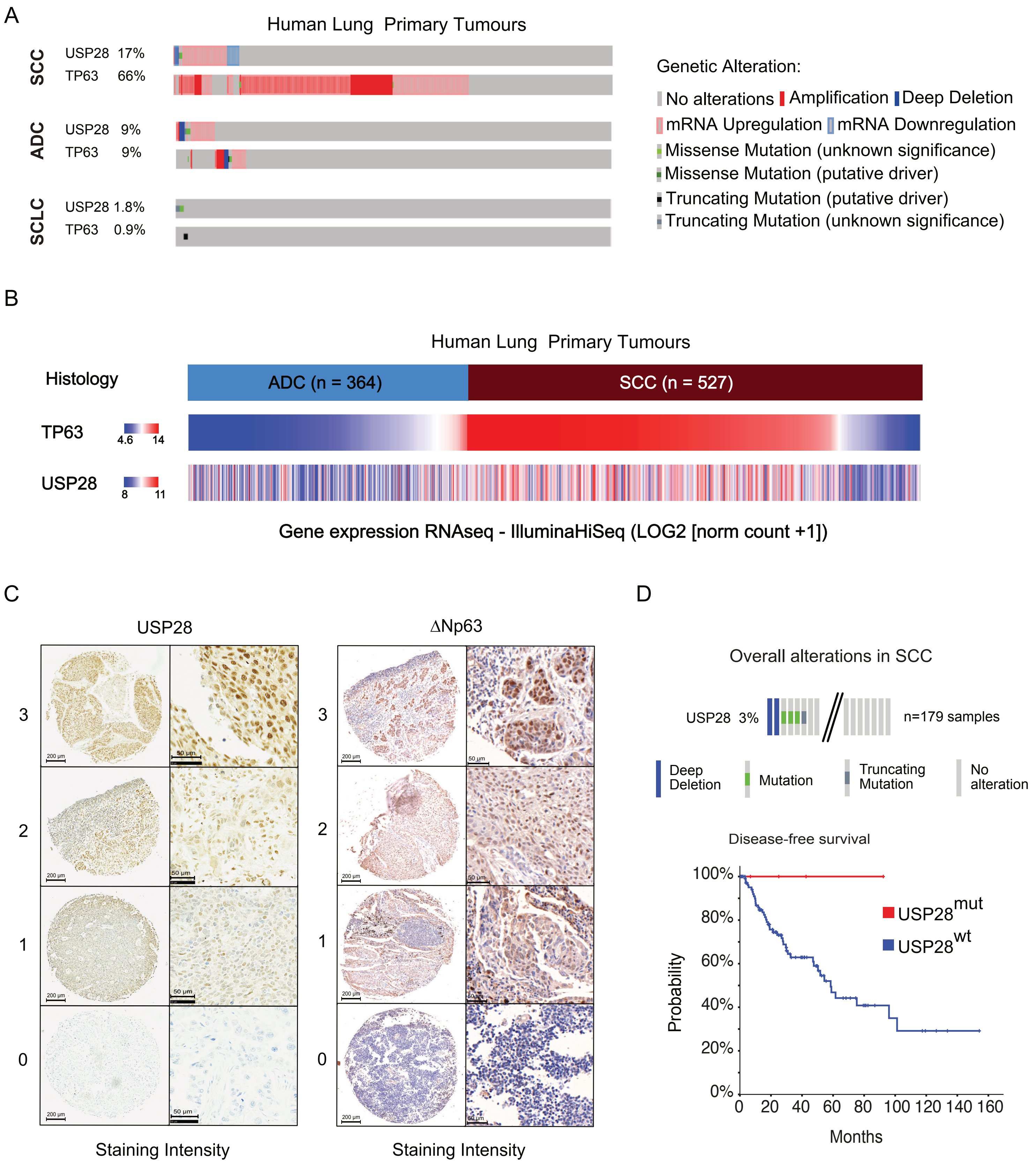


**Supplementary Figure 1: USP28 and ΔNp63 mRNA and protein expression in public data sets, TMA and patient material.**
A) Analysis of occurring genetic alterations in USP28 and TP63 in lung cancer (www.cbioportal.org), B) USP28 and TP63 genomic signature in ADC (n=364) and SCC (527) lung cancer samples (www.xenabrowser.net) C) Representative IHC grading scores of endogenous USP28 and ΔNp63 in lung tissue samples (left panel, low magnification, right panel high magnification), D) Genetic alterations of USP28 in human lung SCC. Each column represents a tumor sample (n = 179 LSCC). Data from TCGA were analyzed using cBioportal abd Xenabrowser software). Disease free survival of USP28 mutant lung SCC patients. Data from TCGA were analyzed using cBioportal software (www.cbioportal.org).

Prieto-Garcia et al., Supplementary Figure 1

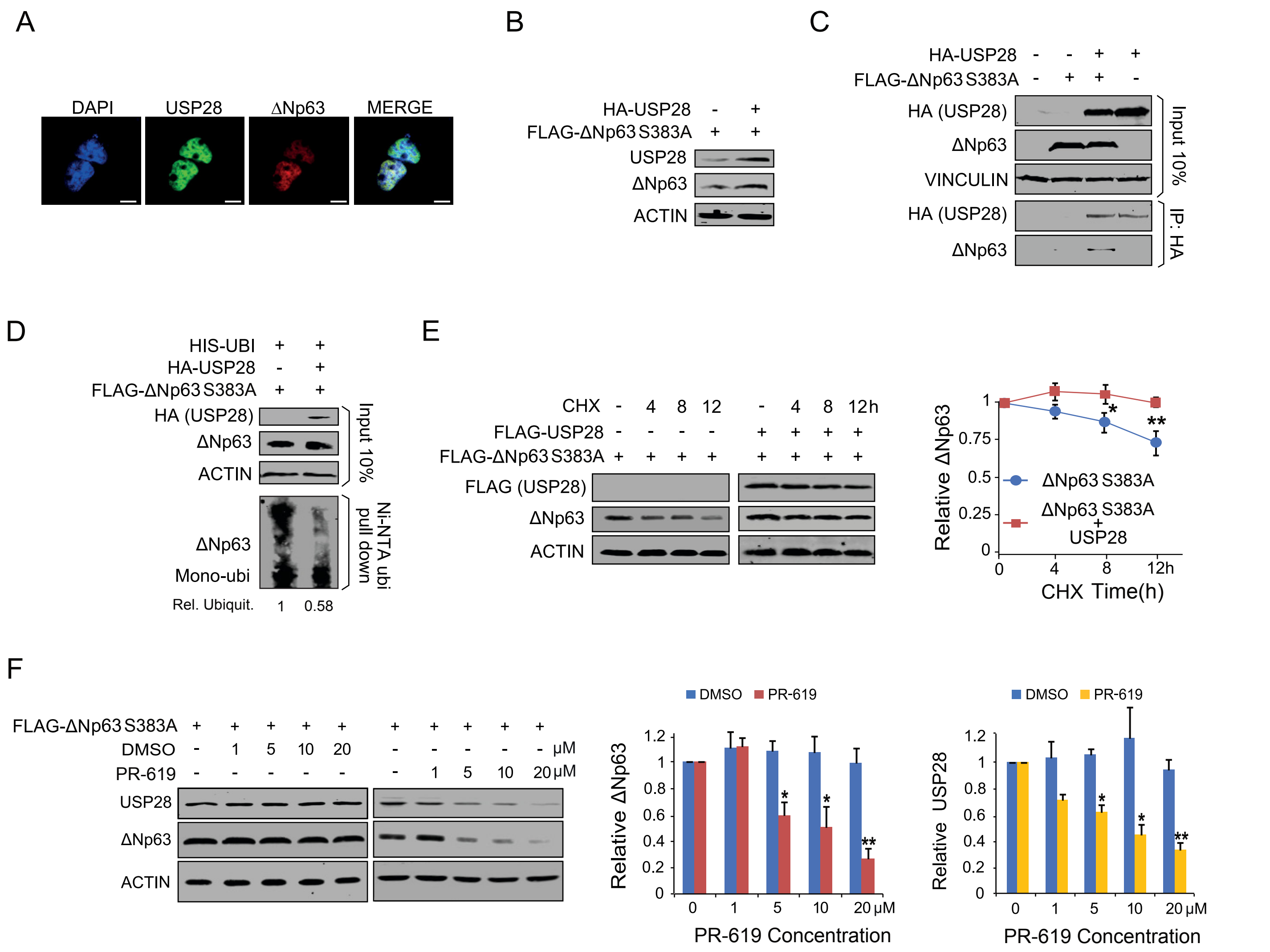


**Supplementary Figure 2: USP28 stabilizes ΔNp63 independently of FBXW7**

A) Immunofluorescence staining against overexpressed HA-USP28 and FLAG-ΔNp63 in HEK293 cells, C) Representative co-immunoprecipitation of exogenous HA-USP28 and FLAG-ΔNp63S383A in HEK293 cells. ACTIN served as loading control. HA-USP28 was precipitated and blotted against FLAG-ΔNp63S383A or HA-USP28 (n=3), D) Ni-NTA His-ubiquitin pulldown of exogenous FLAG-ΔNp63S383A in control transfected or HA-USP28 overexpressing HEK293 cells, followed by immunoblot against ΔNp63S383A protein (Relative Ubiquitination calculated using ACTIN for normalization); E) Immunoblot of HEK293 cells transiently co-transfected with FLAG-ΔNp63S383A together with FLAG-USP28, followed by CHX chase for indicated timepoints. ACTIN served as loading control. Quantification of relative protein abundance of three individual experiments; F) Immunoblot of HEK293 cells transiently co-transfected with FLAG-ΔNp63S383A. 24 hours later cells were treated with either DMSO or the pan-DUB inhibitor PR619 for 24 hours for indicated concentrations., followed by immunoblot against endogenous USP28 and FLAG-ΔNp63S383A. ACTIN served as loading control. Quantification of relative protein abundance of three individual experiments.

All quantitative data are represented as mean ± SD; *$p < 0.05$, **$p < 0.01$

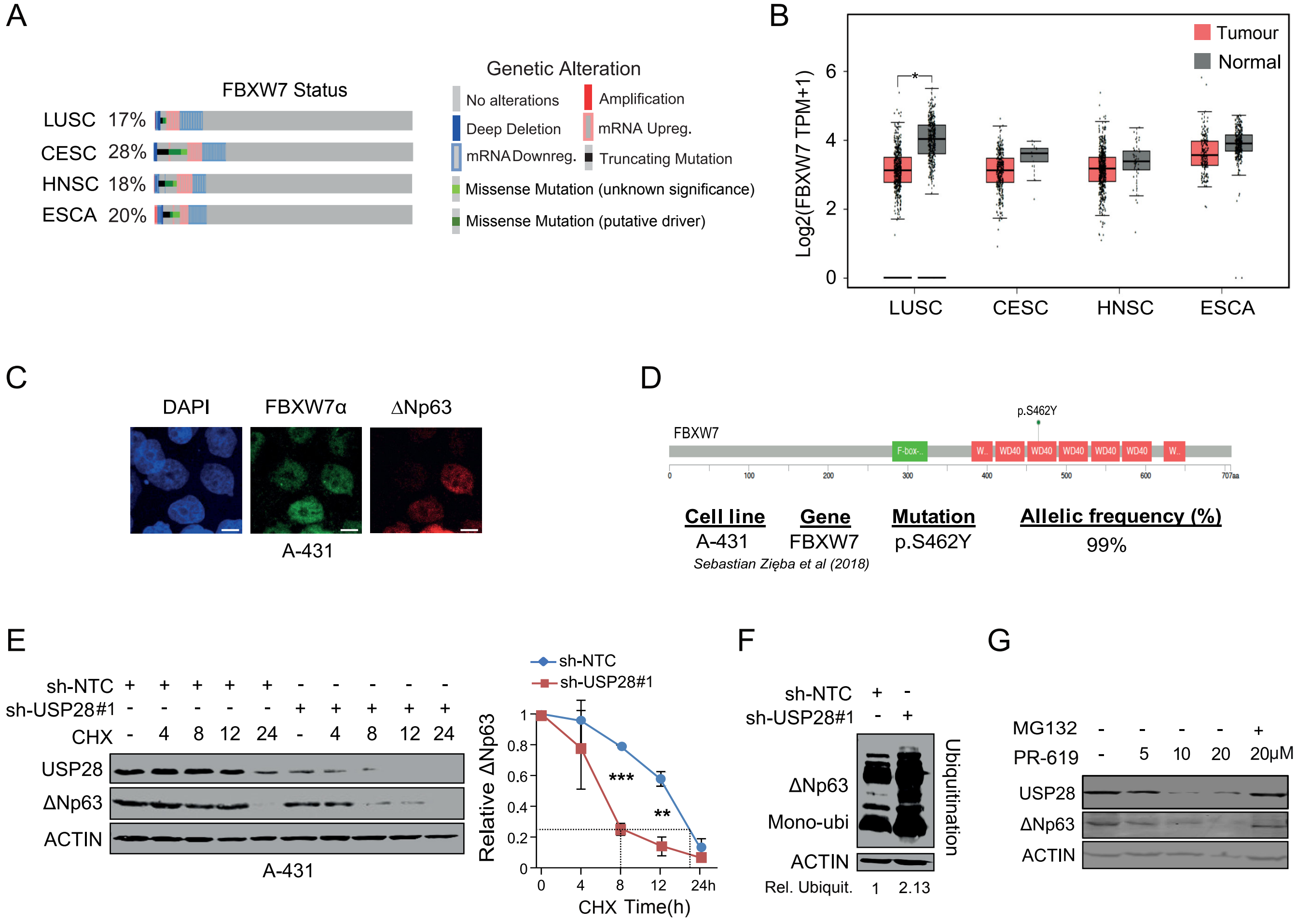


**Supplementary Figure 3: USP28 modulates ΔNp63 proteasomal degradation in FBXW7 deficient cells**
A) Analysis of occurring genetic alterations of FBXW7 in lung-, cervical-, head and neck- and oesophageal squamous tumours (www.cbioportal.org); B) Relative mRNA expression data of FBXW7 in lung-, cervical, head and neck- and oesophageal carcinomas compared to non-transformed tissue samples (www.gepia.cancer-pku.cn); C) Immunofluorescence staining against endogenous FBXW7 and ΔNp63 in the human SCC cell line A431; D) mutational analysis of FBXW7 in A431 (www.cancer.sanger.ac.uk/cosmic); E) CHX chase for indicated timepoints, followed by Immunoblot against endogenous USP28 and ΔNp63, in A431 cells stably transduced with constitutive shRNA-non-targeting control (NTC) or against USP28. VINCULIN served as loading control. Quantification of protein stability of ΔNp63 in three individual experiments; F) TUBE ubiquitin pulldown against endogenously ubiquitylated ΔNp63 in constitutive shRNA-NTC or USP28 knock down A431 cells, followed by immunoblot against ΔNp63. ACTIN served as loading control to calculate relative ubiquitination. G) Treatment of A431 cells with the pan-DUB inhibitor PR-619 for 24 hours at indicated time-points, followed by MG132 freatment for 6 hours at the highest PR-619 concentration, followed by Immunoblot against endogenous USP28 and ΔNp63. ACTIN served as loading control.
All quantitative data are represented as mean ± SD; *$p < 0.05$, **$p < 0.01$; *** $p < 0.001$

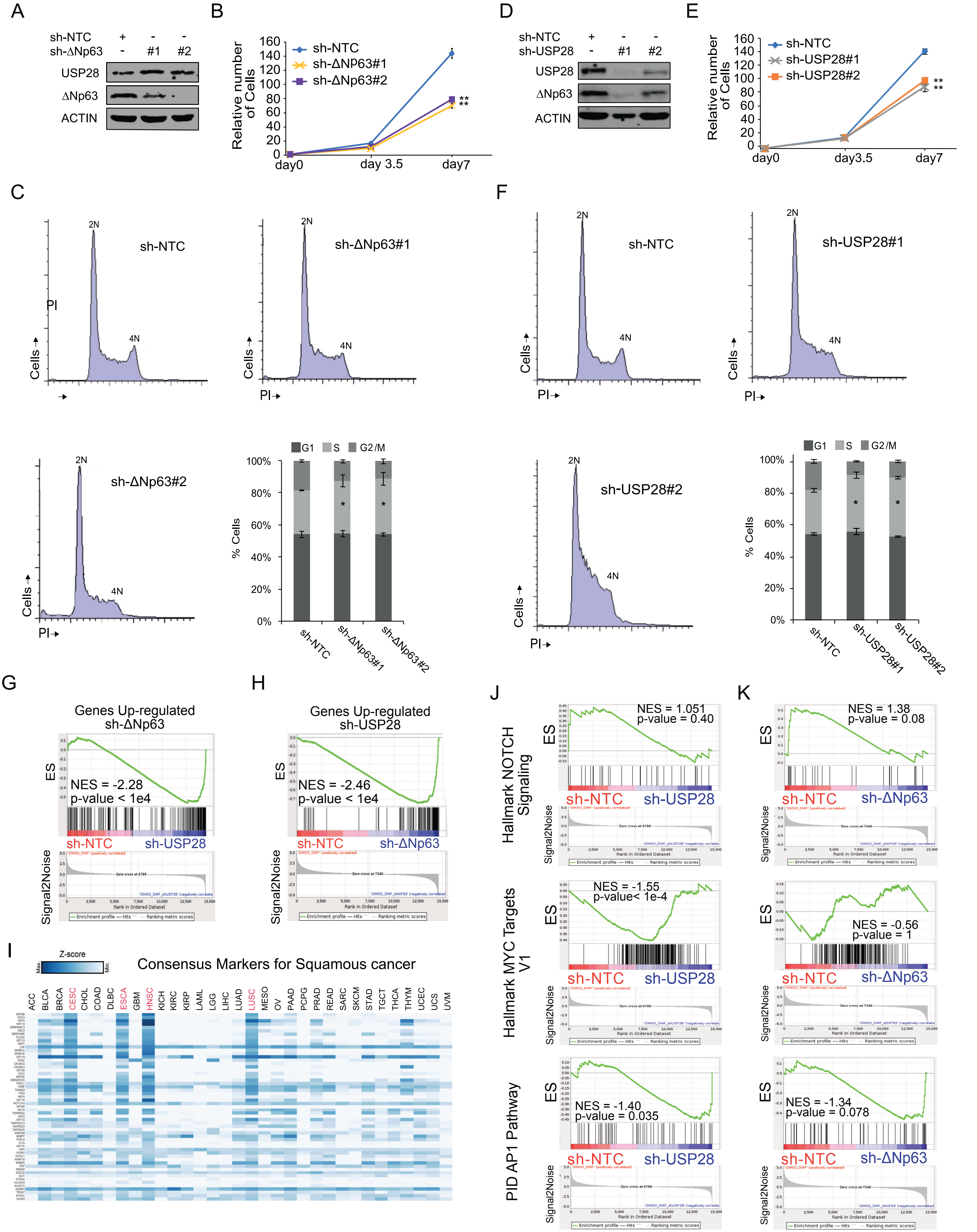


Prieto-Garcia et al., Supplementary Figure 4

**Supplementary Figure 4: SCC tumour cells are dependent on USP28 and/or ΔNp63 to maintain a SCC identity**

A) Immunoblot of endogenous ΔNp63 and USP28 in A431 cells stably transduced with shRNA-non-targeting control (NTC) and two sh-RNA against ΔNp63. Actin served as loading control. B) Cell growth of A431 cells stably transduced with shRNA-non-targeting control (NTC) and two sh-RNA against ΔNp63. Total cell number was measured in triplicate and assessed at indicated time points. C) Cell cycle profile analysis by propidium iodide staining of stable ΔNp63 knock down A431 cells by two independent shRNA sequences. D) Immunoblot of endogenous ΔNp63 and USP28 in A431 cells stably transduced with shRNA-non-targeting control (NTC) and two sh-RNA against USP28. Actin served as loading control. E) Cell growth of A431 cells stably transduced with shRNA-non-targeting control (NTC) and two sh-RNA against USP28. Total cell number was measured in triplicate and assessed at indicated time points. F)Cell cycle profile analysis by propidium iodide staining of stable USP28 knock down A431 cells by two independent shRNA sequences. G) Gene set enrichment analyses of USP28#1 silenced A431 cells compared to shRNA-NTC using the gene list: "Genes Up-regulated sh-ΔNp63". NES, normalized enrichment score; $p < 1e4$. H) Gene set enrichment analyses of ΔNp63 silenced A431 cells compared to shRNA-NTC using the gene list: "Genes Up-regulated sh-USP28". NES, normalized enrichment score; $p < 1e4$Gene. I) Relative expression of consensus markers for Squamous Cancer, as used in Figure3 H and I, in a pan-cancer panel (www.gepia.cancer-pku.cn). J) Gene set enrichment analyses of USP28 silenced A431 cells compared to shRNA-NTC using the gene list: "Hallmark NOTCH Signaling", "Hallmark MYC targets V1' and "PID AP1 Pathway". NES, normalized enrichment score. K) Gene set enrichment analyses of ΔNp63 silenced A431 cells compared to shRNA-NTC using the gene list: "Hallmark NOTCH Signaling", "Hallmark MYC targets V1' and "PID AP1 Pathway". NES, normalized enrichment score

All quantitative data are represented as mean ± SD; *$p < 0.05$, **$p < 0.01$.

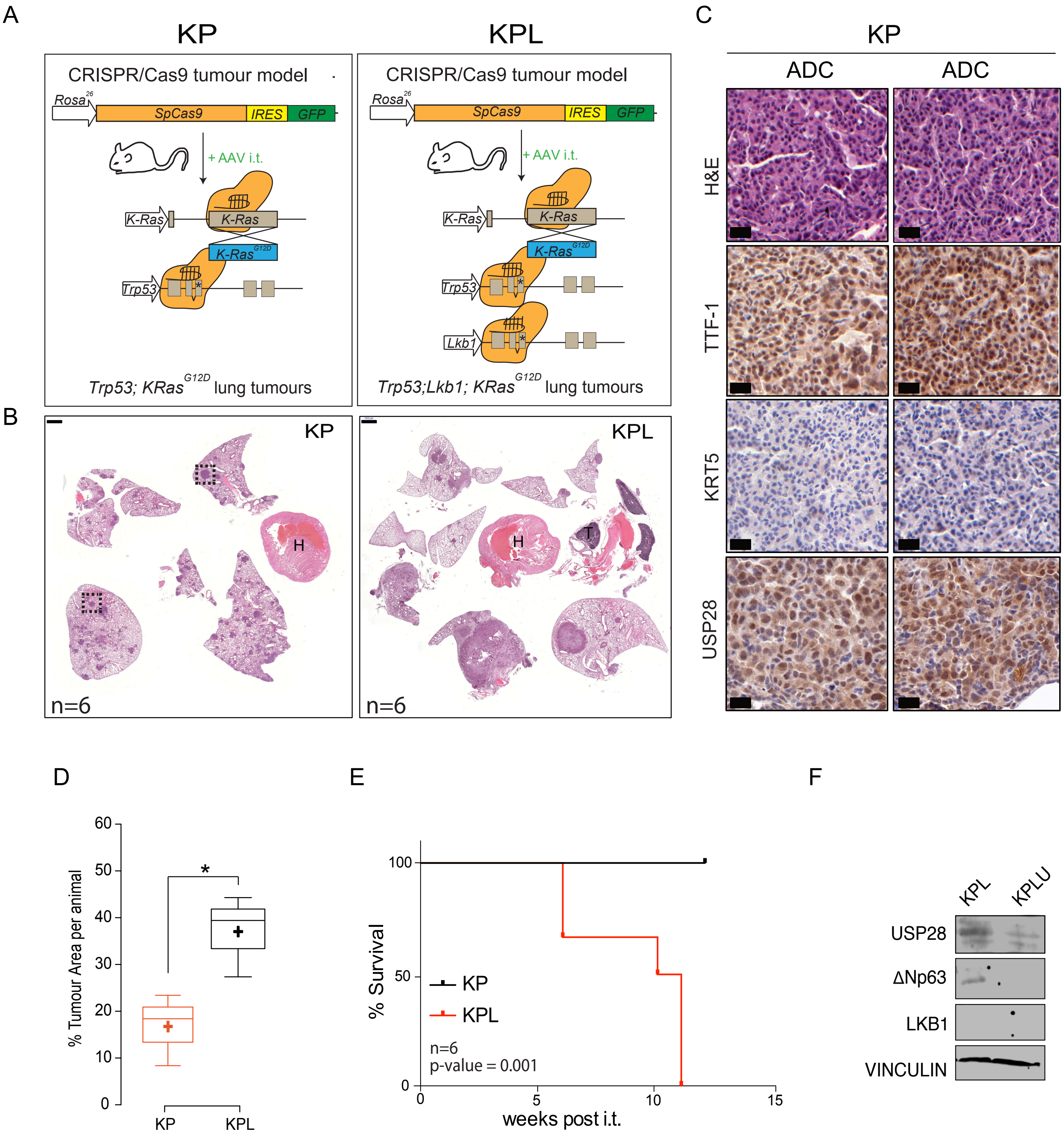


**Supplementary Figure 5: Loss of Lkb1 induces SCC tumour formation, reduces lifespan and Usp28 is upregulated in SCC.** A) Schematic diagram of CRISPR/Cas9 mediated tumour modelling and targeting of $p53^{\Delta}$ : $KRas^{G12D}(KP^{cc})$ or $p53^{\Delta}$; $Lkb1^{\Delta}$: $KRas^{G12D}(KPL^{cc})$ mouse lines; B) Representative H&E images of tumour bearing animals 12 weeks post intratracheal infection. Boxes indicate individual tumour areas assessed by IHC against marker proteins and Usp28 (H= heart, T= thymus, scale bar: 1000um); C) IHC analysis of ADC and SCC marker expression, as well as Usp28 abundance, in KP and KPL lung tumours (scale bar: 20um); D) Boxplot analysis of % tumour area and tumour numbers, per animal, in KP and KPL animals; E) Kaplan Meyer blot of comparing KP versus KPL animals (Mantel Cox test, p = 0.001; n=6). F) Immunoblot of endogenous ΔNp63, USP28 and LKB1 in KPL (SCC) and KPLU (VINCULIN as a control).
All quantitative data are represented as mean ± SD; $*p < 0.05$, $**p < 0.01$

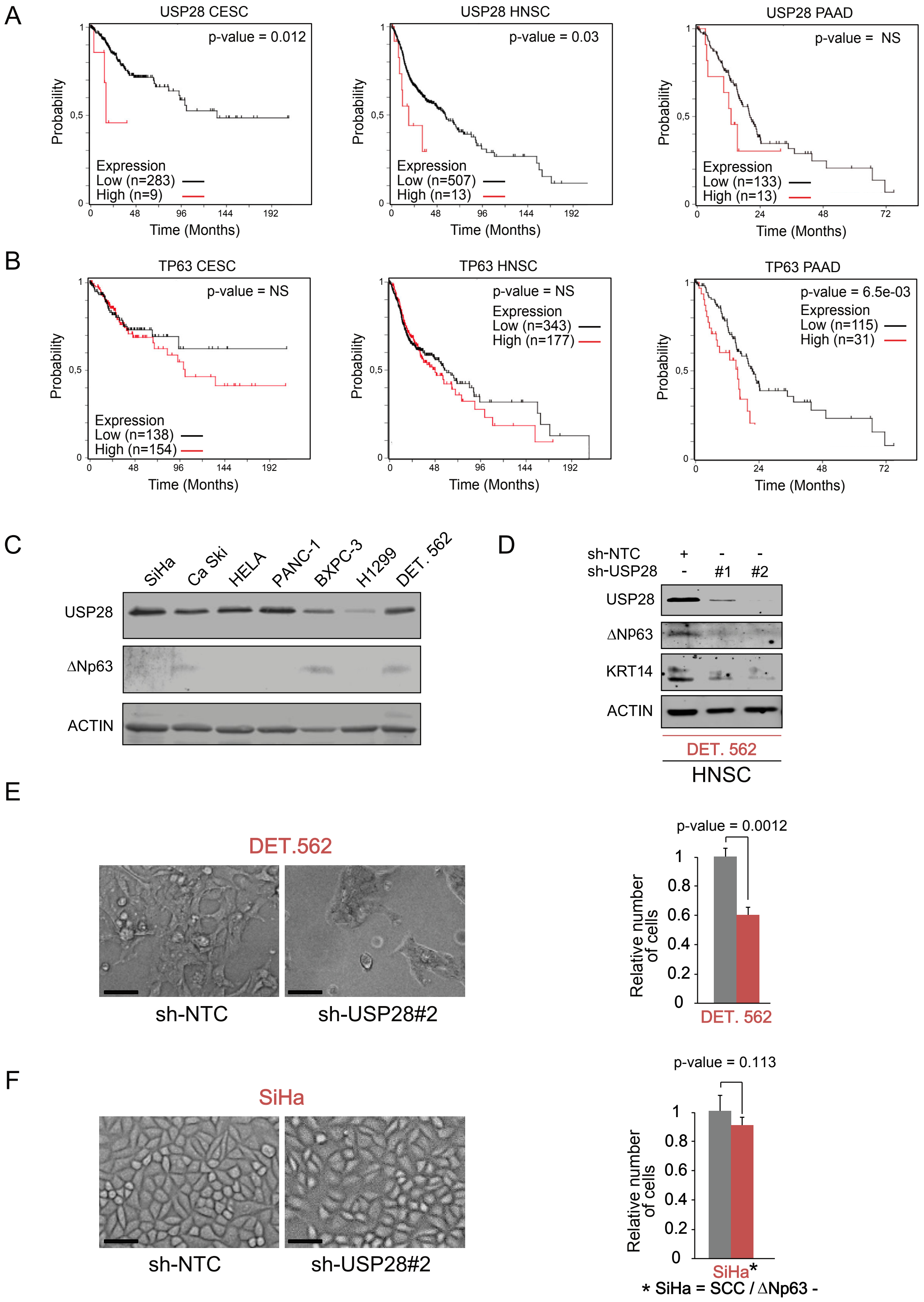


**Supplementary Figure 7: ΔNp63-driven SCC cells of various tissues are vulnerable to USP28 depletion.**
A) Kaplan-Meier estimator of patients survival in cervical SCC (CESC, n=292), head-and-neck SCC (HNSC, n=520) and pancreatic adenocarcinoma (PAAD, n=146) by USP28 expression. p-values were calculated using log-rank test. B) Kaplan-Meier estimator of patients survival in cervical SCC (CESC, n=292), head-and-neck SCC (HNSC, n=520) and pancreatic adenocarcinoma (PAAD, n=146) by ΔNp63 expression. Notably, only PAAD was significant for ΔNp63, indicating the SCC subtype. p-values were calculated using log-rank test. C) Immunoblot against endogenous USP28 and ΔNp63 in human CESC, PAAD, lung ADC and HNSC. Actin served as loading control. Notably, the human CESC cell line SiHa was negative for ΔNp63. D) Immunoblot of USP28, ΔNp63 and KRT14 in HNSC cell line Detroit 562 transduced with non-targeting (sh-NTC) or two independent shRNA against USP28 (shUSP28#1 and #2). ACTIN served as loading control. E) Brigthfield images of Detroit 562 were stably transduced with sh-NTC or shUSP28#2, seeded at equal cell density and counted after five days. Relative number of sh-USP28#2 cells compared with the sh-NTC control cells. p-values were calculated using two-tailed t-test statistical analysis. Scale bar = 30µm F) Brigthfield images of SiHa were stably transduced with sh-NTC or shUSP28#2, seeded at equal cell density and counted after five days. Relative number of sh-USP28#2 cells compared with the sh-NTC control cells. p-values were calculated using two-tailed t-test statistical analysis. Scale bar = 30µm
