## Supplementary material for "The USP28-ΔNp63 axis is a vulnerability of squamous tumours": Consensus Squamous Marker Signature Genes

**Squamous Markers**

ALOX15

ALOXE3

CALML3

CALML5

CLCA2

DSC3

DSG3

EPHA1

EPHA5

FAT2

FGFR2

FGFR3

FREM2

GLI1

IGFBP3

IL1A

IRF5

JUP

Klk10

KRT13

KRT13

KRT14

KRT15

KRT16

KRT19

KRT3

KRT34

KRT4

KRT5

KRT6A

KRT6B

KRT6C

KRT75

KRT84

MIR205HG

MMP10

MMP28

MMP7

NOTCH3

OVOL1

PERP

PGF

PKP1

PVRL1

PVRL4

RAB25

SERPINB13

SERPINB5

SOCS2

SOX2

SPRR1A

SPRR1B

TMPRSS13

TMPRSS3

TMPRSS4

TMPRSS5

TP63

TP63

TRIM29

TRIM7
